# High-throughput Identification of Novel Heat Tolerance Genes via Genome-wide Pooled Mutant Screens in the Model Green Alga *Chlamydomonas reinhardtii*

**DOI:** 10.1101/2022.07.13.499508

**Authors:** Erin M. Mattoon, William McHargue, Catherine E. Bailey, Ningning Zhang, Chen Chen, James Eckhardt, Chris G. Daum, Matt Zane, Christa Pennacchio, Jeremy Schmutz, Ronan C. O’Malley, Jianlin Cheng, Ru Zhang

**Author notes:** Corresponding author: Ru Zhang.

## Abstract

Different high temperatures adversely affect crop and algal yields with various responses in photosynthetic cells. The list of genes required for thermotolerance remains elusive. Additionally, it is unclear how carbon source availability affects heat responses in plants and algae. We utilized the insertional, indexed, genome-saturating mutant library of the unicellular, eukaryotic green alga *Chlamydomonas reinhardtii* to perform genome-wide, quantitative, pooled screens under moderate (35°C) or acute (40°C) high temperatures with or without organic carbon sources. We identified heat-sensitive mutants based on quantitative growth rates and identified putative heat tolerance genes (HTGs). By triangulating HTGs with heat-induced transcripts or proteins in wildtype cultures and MapMan functional annotations, we present a high/medium-confidence list of 933 *Chlamydomonas* genes with putative roles in heat tolerance. Triangulated HTGs include those with known thermotolerance roles and novel genes with little or no functional annotation. About 50% of these high-confidence HTGs in Chlamydomonas have orthologs in green lineage organisms, including crop species. *Arabidopsis thaliana* mutants deficient in the ortholog of a high-confidence *Chlamydomonas* HTG were also heat sensitive. This work expands our knowledge of heat responses in photosynthetic cells and provides engineering targets to improve thermotolerance in algae and crops.

## Introduction

Global warming increases the frequency of damagingly high temperatures, which jeopardize plant growth and reduce food/biofuel production (Janni *et al*. 2020). Nine of the ten hottest years on record have occurred between 2010-2021 (NASA, Goddard Institute for Space Studies). Global temperatures are predicted to reach 1.5°C above pre-industrial levels between 2030 and 2052 at the current warming rate (Intergovernmental Panel on Climate Change, IPCC, 2018) (Jagadish, Pal, Sukumaran, Parani & Siddique 2020). It is estimated that every degree Celsius increase in mean global temperature reduces global yields of wheat, rice, maize and soybean by 6%, 3.2%, 7.4% and 3.1%, respectively (Zhao *et al*. 2017). A recent model based on weather information and six major US crops from 1981-2017 identified high temperatures as the primary climatic driver for yield reduction in crops (Ortiz-Bobea, Wang, Carrillo & Ault 2019). It is essential to understand how plants respond to high temperatures and which genes are required for thermotolerance to engineer heat tolerant crops for the increasing human population.

High temperatures affect many cellular processes in plants (Mittler, Finka & Goloubinoff 2012; Schroda, Hemme & Mühlhaus 2015; Janni *et al*. 2020). They increase membrane fluidity (Martinière *et al*. 2011), ion channel activities, and intracellular calcium concentrations (Saidi *et al*. 2009; Mittler *et al*. 2012), reduce photosynthesis (Zhang & Sharkey 2009; Anderson *et al*. 2021; Zhang *et al*. 2022a), cause protein unfolding/misfolding and modifications (Scharf & Nover 1982; Duncan & Hershey 1989; Schroda *et al*. 2015; Rütgers *et al*. 2017; Wang *et al*. 2020), alter cellular metabolites (Hemme *et al*. 2014; Rysiak *et al*. 2021; Jamloki, Bhattacharyya, Nautiyal & Patni 2021), and affect DNA/RNA stability in the nucleus (Kantidze, Velichko, Luzhin & Razin 2016; Su *et al*. 2018). High temperatures also increase reactive oxygen species (ROS) production from chloroplasts and mitochondria (Pospíšil 2016; Janni *et al*. 2020; Niemeyer, Scheuring, Oestreicher, Morgan & Schroda 2021). Additionally, high temperatures induce heat shock transcription factors (HSFs) which increase the expression of heat shock proteins (HSPs) and other heat response genes (Schroda *et al*. 2015; Guo *et al*. 2016).

Several major questions remain open regarding heat responses in plants, despite our current understanding as mentioned above (Mittler *et al*. 2012; Schroda *et al*. 2015; Vu, Gevaert & De Smet 2019). (1) What are the genes required for heat tolerance? Even with a partial list of such genes, we can better understand heat tolerance in photosynthetic cells and provide engineering targets to make heat tolerant crops. (2) Do different high temperatures require distinct or overlapping genes for heat tolerance? High temperatures have different intensities in nature, which have unique effects on photosynthetic cells (Janni *et al*. 2020; Zhang *et al*. 2022a). Moderate high temperature refers to heat that is slightly above optimal growth temperature, around 35°C for most photosynthetic cells, which causes moderate damages and occurs frequently with prolonged duration in nature (Zhang *et al*. 2022a). Acute high temperature refers to heat at or above 40°C, which occurs less frequently but can cause severe damage to cellular processes (Zhang *et al*. 2022a). (3) How does carbon supply affect heat tolerance? Land plants mostly grow photoautotrophically by fixing atmospheric inorganic carbon via photosynthesis, however, there are often various organic carbon sources in the soil. Some plant tissues are mixotrophic (e.g., seed pods) (Koley *et al*. 2022), partially replying on transported carbon sources from leaves. Most green algae can grow both photoautotrophically and mixotrophically with supplied organic carbon. How carbon supply affects heat responses in photosynthetic cells is understudied.

The unicellular green alga *Chlamydomonas reinhardtii* (Chlamydomonas throughout) is a powerful model to study heat responses in photosynthetic cells. It has a sequenced, haploid, and simple genome (111 Mb, 17,741 protein-encoding genes) with smaller gene families and lower rates of gene duplication compared to land plants, simplifying genetic analyses (Merchant *et al*. 2007; Karpowicz, Prochnik, Grossman & Merchant 2011). It grows fast with a 6-8 h doubling time under normal conditions. It can grow solely by photosynthesis in light or with a supplied carbon source (acetate) in light or dark, convenient for laboratory experiments and strain maintenance (Harris 2009). Photosynthesis is one of the most heat sensitive functions in plants (Sharkey 2005; Sharkey & Zhang 2010). The photosynthesis in Chlamydomonas is highly similar to land plants (Minagawa & Tokutsu 2015), and its ability to survive on organic carbon enables the maintenance of photosynthetic mutants, making it a superior model to study heat effects on photosynthesis. Additionally, several excellent molecular and genetic tools are available for Chlamydomonas, enabling efficient gene editing (Shimogawara, Fujiwara, Grossman & Usuda 1998; Greiner *et al*. 2017; Wang *et al*. 2019; Dhokane, Bhadra & Dasgupta 2020) and molecular engineering (Crozet *et al*. 2018; Emrich-Mills *et al*. 2021). Specifically, a mapped, indexed, genome-saturating Chlamydomonas insertional mutant library is available for both reverse and forward genetic screens (Chlamydomonas Library Project, CLiP, 62,389 mutants, covering 83% of nuclear protein-coding genes) (Zhang *et al*. 2014, 2022b).

Each CLiP mutant contains at least one unique DNA barcode by the end of the insertional cassette, allowing for high-throughput, quantitative tracking of cell abundance via deep sequencing and growth rate calculation of each individual mutant in pooled cultures under different conditions (Li *et al*. 2016, 2019). If a CLiP mutant is deficient in a gene required for optimal growth under a condition, this CLiP mutant will have a reduced cell abundance, barcode abundance, and growth rate under this condition. The unique DNA barcode is also linked to a mapped insertion site in each mutant, allowing for identification of genes important for growth under a defined condition.

The Jonikas laboratory, which led the generation of the CLiP mutant library, together with the Jinkerson/Dinneny groups and other collaborators, employed the CLiP library to screen for mutants under 121 different environmental/chemical conditions (Fauser *et al*. 2022). The pooled screens by Fauser et al. (2022) have been foundational in using the CLiP mutant library for functional genomics. They utilized a broad range of experimental conditions for functional prediction of genes with unknown roles and this work has been a pivotal advancement in the field. Among 121 conditions, Fauser et al. (2022) included but did not focus on high temperatures. They included high temperatures of 30°C, 35°C, 37°C and tested the effects of light intensities, CO_2_ concentration, and organic carbon availability on heat responses (Supplemental Dataset 1a). The high-temperature screens by Fauser et al. (2022) had several limitations. (1) They emphasized 30°C heat which is a rather mild high temperature for CC-5325, the background strain of the CLiP mutant library (Zhang *et al*. 2014, 2022), and did not include heat at 40°C, which is acute high temperature for CC-5325. (2) They did not include heat at 35°C without carbon supply for parallel comparison with carbon supply at 35°C. (3) Their heat screens were performed in 2-L medium bottles on heat stirring plates with constant setting-temperature but without control of culture temperatures, cell density, and nutrient supplies, which may complicate the screen results. (4) The CLiP mutants were streaked off the algal plates, inoculated into liquid cultures, and grown under different temperatures directly, without solid to liquid acclimation process before heat treatments, which may increase the false positive rates of heat-sensitive mutants. (5) Due to the large-scale nature of their experimental setup, they had single replicates for 6 out of 11 different heat conditions performed. (6) They did not have transcriptomes or proteomes under high temperatures to refine their candidate genes identified in the heat screens.

Our research focused on high temperature conditions using the CLiP mutant library and the quantitative pooled screens. We advanced the high temperature screens by Fauser et al. (2022) to identify more genes important for heat tolerance. Based on the growth rates of the CLiP library background strain, CC-5325, under different temperatures, we performed pooled CLiP mutant screens under moderate (35°C) and acute high temperature (40°C) with and without organic carbon supply in photobioreactors (PBRs) under well-controlled conditions. Our algal cultivation in PBRs had turbidostatic control, which allowed for precise control of growth environments including temperature regulation, heating speed, light intensity, air bubbling, cell density, and, importantly, nutrient supply. In our cultivation method, we can isolate the effects of high temperatures on algal cells and minimize the complications from other environmental factors, such as nutrient depletion or light shading. Furthermore, we developed a triangulation approach to combine CLiP pooled screens, our recently published wildtype (WT) Chlamydomonas transcriptome and proteome data under 35°C and 40°C, and functional annotations to identify triangulated, known and novel, heat tolerance genes (HTGs). We further sorted these triangulated HTGs into high/medium-confidence levels based on the presence of multiple heat-sensitive alleles. Additionally, our orthology analysis revealed that many of the triangulated HTGs identified in Chlamydomonas are conserved in land plants. Finally, *Arabidopsis thaliana* (Arabidopsis throughout) mutants deficient in the orthologous gene of a Chlamydomonas high-confidence HTG were heat sensitive. Our research provides engineering targets to improve thermotolerance in both green algae and land plants.

## Materials and Methods

### Algal cultivation

All Chlamydomonas liquid cultivation used in this paper were conducted in photobioreactors (PBRs) as described before (Zhang *et al*. 2022a) with minor modifications. Algal cultures were grown in standard Tris-acetate-phosphate (TAP, with acetate, an organic carbon source) or Tris-phosphate (TP, without acetate) medium with modified trace elements (Kropat *et al*. 2011) in 1-L PBRs (Photon System Instruments, FMT 150/1000-RB). Cultures were illuminated with constant 100 µmol photons m^2^ s^−1^ light (50% red: 50% blue) and mixed by bubbling with filtered air. After inoculation, cultures grew to a target cell density of 2×10^6^ cells/mL in log-phase growth at 25°C. Then, the target cell density was maintained turbidostatically using OD_680_ (monitored every 1 min automatically) by allowing the culture to grow to 8% above the target cell density before being diluted to 8% below the target cell density with fresh medium provided through peristaltic pumps. Through the turbidostatic mode, the PBR cultures had constant nutrient supply, controlled cell density, and exponential growth between dilution events. The OD_680_ measurements during exponential growth phases in between dilution events were log_2_ transformed, and the relative growth rate was calculated using the slope of log_2_(OD_680_) while the inverse of the slope yielded the doubling time of the culture (Zhang *et al*. 2022a). OD_680_ is a non-disruptive, efficient, and automatic measurement with 1-min resolution and high sensitivity, which is well suited for algal growth measurement under our cultivation conditions with turbidostatic control and constant dilutions. We used a small OD_680_ range for turbidostatic control. OD_680_ data is sensitive enough to quantify algal growth during each short exponential growth phase. The growth rates calculated by OD_680_ was mostly for us to estimate the stress level of different high temperature treatments. Different treatments (different temperatures and medium type) were conducted in individual PBRs. For WT CC-5325 cultures, cells were first acclimated at 25°C for 2-days until reaching steady growth rates before treatments at different high temperatures. Each high temperature treatment was conducted in individual PBRs with replicates.

For pooled screens, the CLiP mutant library was pooled as described before (Fauser *et al*. 2022) with minor modifications. The CLiP library was replicated using a Singer RoToR robot (Singer Instruments, 704) on 384-format 1.5% TAP agar plates and grown in the dark. Five-day-old fresh plates with the CLiP mutant library were pooled using sterile glass spreaders in liquid TAP medium, vortexed to break the colonies, grown on a shaker in the dark at room temperature overnight before being inoculated into PBRs the next day. This gave cells time to acclimate from solid agar to liquid cultures before inoculation to PBRs. Pooled CLiP liquid cultures were diluted using TAP or TP medium before being inoculated into PBRs at 25°C in TAP or TP medium (initial cell density around 1×10^6^ cells mL^-1^) and acclimated at 25°C in the light for 2 days to reach steady growth rates with turbidostatic control before the start of heat treatments. Acclimation at 25°C in the light was used to reduce false-positive heat-sensitive mutants that have difficulty adjusting from growth on solid to liquid medium or from dark to light conditions. The heat of 35°C lasted 4 (TAP) or 5 (TP) days and the heat of 40°C lasted 2 days because the algal cultures would not survive if heated at 40°C for longer. After 40°C heat, cultures recovered at 25°C for 3 days. Algal cultures of pooled mutants (2 x 50 mL) were harvested before and after the heat treatments or by the end of recovery using centrifugation at 4°C for 5 min. Cell pellets were stored at −80°C before DNA extraction and barcode amplification.

### DNA extraction and sequencing

Genomic DNA was extracted for DNA barcode amplification as described before (Li *et al*. 2016) with minor modifications. Frozen cell pellets (from one copy of 50 mL cultures) were mixed with 800 µL of SDS-EDTA buffer (1% SDS, 200 mM NaCl, 20 mM EDTA, 50 mM Tris-HCL, pH 8) and separated into two equal aliquots for downstream processing. Then, 500 µL of 24:25:1 phenol:chloroform:isoamyl alcohol (Sigma, P2069-400ML) was added to each tube. Samples were vortexed for 2 min, centrifuged for 5 min at 10,000 g, and the aqueous phase was aliquoted into a new PhaseLock tube (VWR, Cat No. 10847-800), added with 1.6 µL of 100 mg/µL RNaseA (ThermoFisher Scientific, Cat. No. 1209102), and incubated at 37°C for 30 min. The phenol/chloroform DNA extraction was repeated three additional times. After the final extraction, samples were transferred to a new 1.5 mL tube and 2.5 volumes cold 100% ethanol was added. Samples were incubated in a - 20°C freezer for 1 h then centrifuged at 16,000 g for 20 min at 4°C. Supernatant was decanted and pellets were washed in 1 mL cold 70% ethanol. Supernatant was decanted, pellets dried for 30 min at room temperature and DNA pellets were resuspended in 50 µL molecular grade water. Like samples were combined into a single tube then quantified by high sensitivity DNA Qubit (ThermoFisher scientific, Cat. No. Q32854).

### DNA barcode amplification and sequencing

DNA barcodes were amplified as described before (Li *et al*. 2016) with minor modifications. Barcodes from the 5’ and 3’ side of the insertional cassette were amplified, sequenced, and processed separately. PCR reactions of 50 µL were prepared as follows: PCR 5’ side barcodes: 17 µL molecular grade water, 10 µL GC buffer, 5 µL DMSO, 1 µL dNTPs at 10 mM, 1 µL MgCl_2_ at 50 mM, 2.5 µL each primer at 10 µM, 1 µL Phusion HotStart polymerase, and 10 µL DNA at 12.5 ng/µL. Samples were amplified using the following PCR cycle for 5’ side barcodes: 98°C for 3 min, 10 cycles of 98°C for 10 s, 58°C for 25 s, 72°C for 15 s, 10 cycles of 98°C for 10 s, 72°C for 40 s. PCR for 3’ side barcodes: 16 µL molecular grade water, 10 µL GC buffer, 5 µL DMSO, 1 µL dNTPs at 10 mM, 2 µL MgCl_2_ at 50 mM, 2.5 µL each primer at 10 µM, 1 µL Phusion HotStart polymerase, and 10 µL DNA at 12.5 ng/µL. Samples were amplified using the following PCR cycle for 3’ side barcodes: 98°C for 3 min, 10 cycles of 98°C for 10 s, 63°C for 25 s, 72°C for 15 s, 9 cycles of 98°C for 10 s, 72°C for 40 s.

PCR products were cleaned and concentrated using the MinElute Gel Extraction kit (Qiagen, Cat. No. 28606). The purified product was separated by gel electrophoresis using a 1.5% agarose gel performed at 130 V for 65 min. PCR bands of 235 bp (5’ side barcodes) and 209 bp (3’ side barcodes) were excised from agarose gel and extracted with the MinElute Gel Extraction kit. Final purified DNA was eluted with 20 µL elution buffer. DNA concentration was quantified using Qubit and the quality of DNA was visualized using 8% TBE gel (Invitrogen, Cat. No. EC62152BOX) performed at 230 V for 30 min. Gel was stained for 5 min with SYBRGold (ThermoFisher Scientific, Cat. No. S11494) prior to image acquisition.

Because samples were pooled together for sequencing, barcode amplification primers contained unique indexes for downstream identification in the data analysis pipeline (Supplemental Dataset 1b). Prior to sequencing, eight DNA samples were pooled together for a total of 23.75 ng of each DNA sample in a final volume of 25 µL. Samples were sequenced at the Department of Energy Joint Genome Institute (DOE JGI) using the Illumina Hi-seq platform. The prepared libraries were quantified using KAPA Biosystems’ next-generation sequencing library qPCR kit and ran on a Roche LightCycler 480 real-time PCR instrument. Sequencing of the flowcell was performed on the Illumina NextSeq500 sequencer using NextSeq500 Mid Output kits, v2, following a 1 x 50 indexed run recipe utilizing a custom sequencing primer. Reads were mapped to each DNA sample using the index code contained in the PCR amplification primer. Each DNA sample had an average of 27 million reads and the minimum number of reads was 8 million.

### Barcode abundance quantification and normalized read count cutoffs

From the Illumina sequencing reads, common cassette sequences were trimmed using Geneious v10.1.3, leaving only the unique DNA barcodes. The number of reads from each unique DNA barcode was quantified, and the total number of reads for each sample was normalized to 100 million. The unique DNA barcodes were mapped to the CLiP mutant library with no mismatch bases allowed. Minimum cutoffs of normalized barcode read count were implemented: ≥150 normalized reads in all T1 samples (beginning of the treatment period for control and the high temperature treatment) and ≥1 read from T3a or T3b (end of the screens) in the control condition (Figure S1). Barcodes from heat-compromised individuals were required to have ≥1 read in T3a or T3b of the high temperature treatments. Barcodes from heat-depleted individuals had 0 reads in at least 1 biological replicate at T3a or T3b of the high temperature treatments.

### Principal component analysis

Principal component analysis was performed on normalized read counts from all barcodes in all biological replicates using the R package FactoMineR (Lê, Josse & Husson 2008).

### Growth rate calculations

The growth rate from each individual barcode was calculated using the following equation:

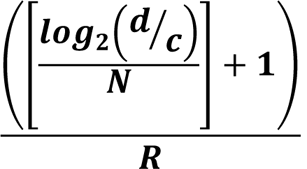

Where d represents the normalized reads at T3a or T3b (end of screens), c represents the normalized reads at T1 (start of screens), and N represents the estimated number of generations that occurred in each given biological replicate (see Supplemental Dataset 1c). These calculations were performed for all barcodes in a biological replicate, then growth rates were divided by R, the mean growth rate of all mutants in the biological replicate to normalize for effects of different treatments. Each mutant was compared to all other mutants in the same biological replicate; thus, the calculated growth rate of each mutant is relative to the whole mutant pool. Relative growth rates were calculated for each barcode in control and high temperature treatments. The mean doubling time of each treatment was calculated based on the exponential increase of OD_680_, as mentioned above. The number of generations that occurred in each biological replicate was estimated by the duration of the experiment divided by the mean doubling time (Supplemental Dataset 1c). The mean growth rate of all mutants in a biological replicate is inverse of the doubling time. The cell cycle was arrested in 40°C (Zhang *et al*. 2022a), thus for 40°C treated cultures, we assumed zero generations occurred during the 2-day 40°C heat, and therefore only estimated the number of generations that occurred during the 25°C recovery period following the 40°C heat treatment. Mutants that were depleted by the end of the 2-day 40°C heat were included as heat-depleted mutants (see below).

### Growth rate validation of individual mutants in monocultures

Ten individual CLiP mutants were selected for monoculture phenotyping. The identities of these mutants were verified by PCR using primers that bind to the cassette or flanking genomic regions (Supplemental Dataset 1b) as described before (Li *et al*. 2016). PCR products were sequenced for confirmation. Mutants were individually grown in PBRs as mentioned above. Cultures were acclimated for 4 days at 25°C, then the temperature was raised to 35°C for 4 days. The exponential increase of OD_680_ was used to determine the doubling time (Td) of the culture, which was then used to calculate the growth rate (1/Td) as described above. For the growth rate at 25°C, doubling times from the last 24 hours of 25°C before heat at 35°C were used to calculate the average pre-heat growth rate. The growth rate at 35°C was calculated from all doubling times from 6 hours after temperature increase to 35°C to the end of 35°C heat. Doubling times from the first 6 hours of 35°C heat treatment tended to be inconsistent as the culture adjusted to the higher temperature. A normalized growth rate was then calculated by dividing the mutant growth rate by the growth rate of WT CC-5325 (CLiP background) for each temperature condition.

### Identification of heat-compromised and heat-depleted mutants

Heat-compromised individuals are required to have a mean normalized growth rate at 25°C ≥0.95 (normal growth) and a mean normalized growth rate at high temperature of ≤0.8 (compromised growth) (Figure S2a-f). Additionally, using the normalized growth rates of two biological replicates at each treatment condition, we performed a student’s one-sided t-test of unequal variance for each barcode that met normalized read count cutoffs. Significance was defined as p < 0.05 and t-value in the 95^th^ percentile (Figure S2g-j). Heat-depleted individuals were also required to have a mean normalized growth rate at 25°C ≥0.95 (normal growth), but because they were completely depleted from at least one biological replicate at high temperature, a growth rate could not be calculated. Heat tolerance genes (HTGs) were defined as those with at least one heat-compromised or heat-depleted mutant. All the pooled screen data with gene IDs, mutant IDs, phenotypes is in Supplemental Dataset 2.

### Comparisons between conditions

To compare the HTGs between conditions, we filtered the dataset for only those genes that were represented by at least one mutant in all four treatment conditions. Comparisons between conditions were visualized using UpSetR (Conway, Lex & Gehlenborg 2017).

### MapMan functional enrichment

Functional enrichment analysis was performed using MapMan for different combinations of conditions including: all HTGs from all conditions, and HTGs overlapping between both TAP conditions, both TP conditions, both 35°C conditions, both 40°C conditions, or all four conditions. Gene lists were expanded such that one gene can be associated with multiple MapMan functional terms. Enrichment was assessed using a Fisher’s Exact Test, FDR < 0.05 (Venn & Mühlhaus 2022; Venn *et al*. 2022; Benjamini & Hochberg 01/1995).

### Transcriptome and proteome comparisons

HTGs from all four conditions were compared with previously published RNA-seq data from WT (CC-1690) Chlamydomonas cells treated for 24 h at either 35°C or 40°C followed by a 48 h recovery period at 25°C in TAP medium (Zhang *et al*. 2022a). Differentially expressed genes were sorted into heat induced genes (HIGs, up-regulated in ≥1 time point during the high temperature period). HIGs were filtered for genes that had ≥1 CLiP mutant present in pooled mutant screens. HTGs were also filtered for those genes that met minimum read count cutoffs in the RNA-seq dataset. Overlaps between HTGs and HIGs, were quantified, and enrichment was determined using a Fisher’s exact test, FDR < 0.05. These RNA-seq data also included modules of genes with similar expression profiles using Weighted Correlation Network Analysis (WGCNA) (Langfelder & Horvath 2008; Zhang *et al*. 2022a). HTGs from the TAP-35°C, TAP-40°C, and the aggregated HTGs from all 4 conditions were sorted within these modules. For modules that contained more HTGs than would be expected by random chance from these groups (Fisher’s exact test, FDR < 0.05), MapMan functional enrichment analysis was performed on the overlapping WGCNA module list and HTG list as described above.

### Heat sensitivity assay of Arabidopsis mutants

Arabidopsis T-DNA insertional mutants were obtained from the Arabidopsis Biological Resource Center (ABRC): *Athmt-1* (SAIL_114_G09) and *Athmt-2* (SALK_106875C), and the positive control *Athot1* (SALK_066374C). Homozygous insertional mutants were verified by PCR as described before (Sessions *et al*. 2002; Alonso *et al*. 2003) (see Supplemental Dataset 1b for primer sequences). WT Col-0 seeds were donated by Dr. Dan Lin at the Donald Danforth Plant Science Center. Seeds from homozygous lines were used for heat sensitivity assays. Seeds were sterilized in 500 µL 20% bleach with 1 µL 20% Tween20 followed by five washes using sterile water, then plated on MS-sucrose plates (0.5X Murashige and Skoog (MS) salts, 1% sucrose), stratified for 3 days in dark at 4°C, then grown in constant white light of 115 µmol photons m^2^ s^−1^ at 25°C in a Conviron growth chamber. After 7 days of growth, 3 biological replicates of seedlings were heated at 41°C for 45 min in a pre-warmed water bath. Heat treated plates were returned to the 25°C growth chamber mentioned above for an additional 7 days for recovery before imaging. Mean plantlet area was measured using ImageJ by dividing the total plantlet area of each genotype by the number of seeds that germinated for that genotype. Mutant plantlet area was normalized to that of Col-0 on the same plate. Statistical significance of mutants compared to WT on the same plates was assessed with a student’s one-sided t-test of unequal variance. Control plates were grown at 25°C in a Conviron growth chamber as mentioned and images were acquired at 10 days of growth for analysis to limit the overlapping of neighboring plants. Gene models for *CrHMT* and *AtHMT* were generated using the Gene Structure Display Server (GSDS2.0) (Hu *et al*. 2015).

## Results

### Pooled mutant screens were conducted at moderate and acute high temperatures

We first grew CC-5325, the background strain of the CLiP mutant library, in photobioreactors (PBRs) under well-controlled conditions to test our heat screen conditions. Cells were grown in PBRs under constant light in either Tris-acetate-phosphate (TAP) medium (contains acetate, the organic carbon source), or Tris-phosphate medium (TP, no acetate) under different temperatures. With supplied carbon, the growth rate of CC-5325 increased at 30°C and 35°C but decreased at 40°C as compared to 25°C (Figure 1a). The increased growth rates at 30°C and 35°C were abolished without supplied organic carbon source, although the cultures were still very sensitive to 40°C. We defined 35°C as moderate high temperature, and 40°C as acute high temperature. We performed genome-wide, quantitative, pooled mutant screens in PBRs using the CLiP mutant library at 25°C, 35°C, and 40°C with or without carbon source (Figure 1b). Algal cultures were harvested at the beginning (T1) and end (T3a or T3b) of the treatment period for DNA extraction, barcode quantification by deep sequencing, and growth rate calculation for each mutant (Figure 1c). Principal component analysis of normalized read abundance from all barcodes in each sample showed high reproducibility between biological replicates (Figure 1d, Figure S1). PC1 explains 40.8% of the dataset variance and separates samples based on medium condition, with and without carbon supply. PC2 explains 11.2% of the dataset variance and separates samples based on both temperature treatments and time points.

**Figure 1:**
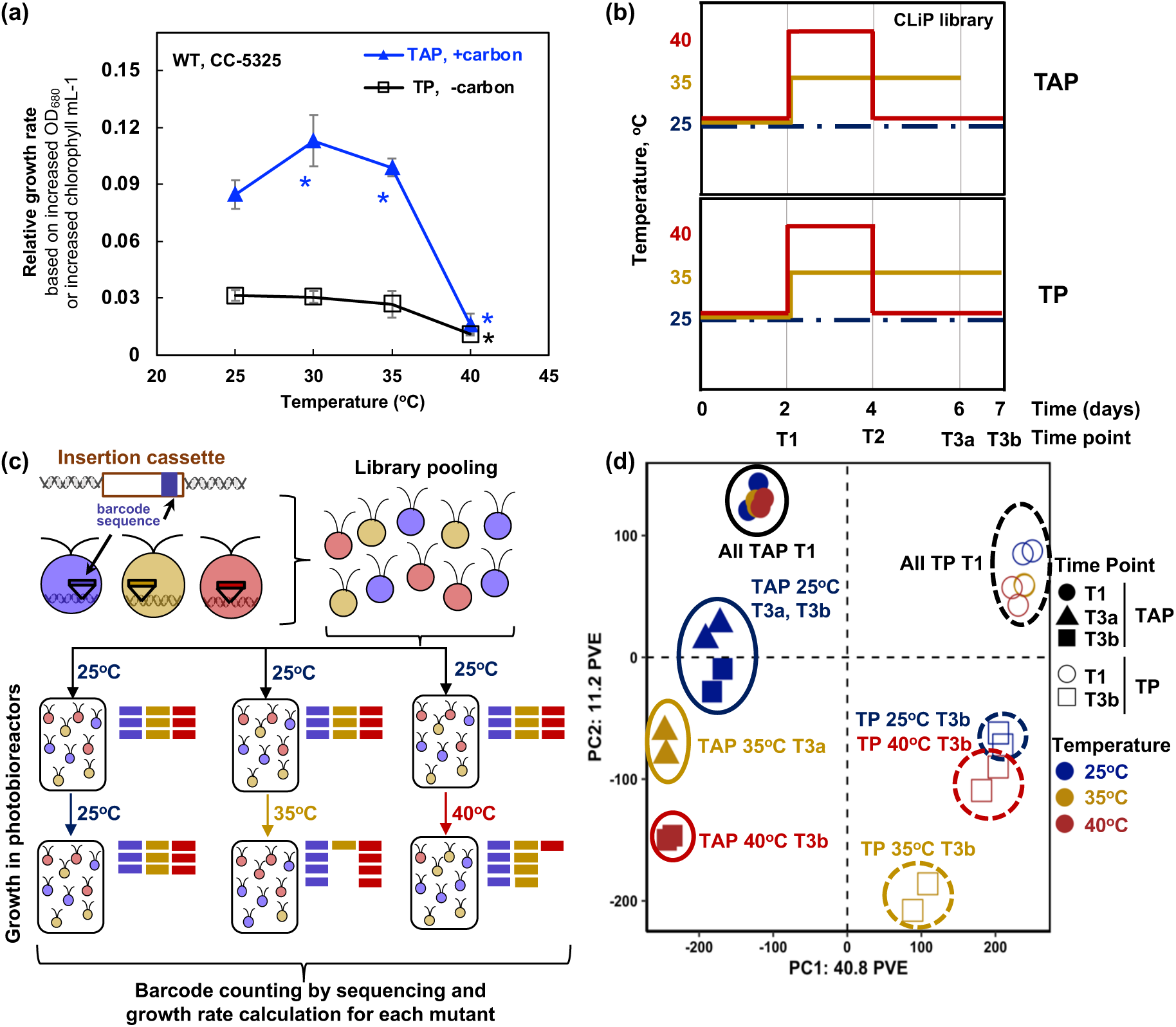
We screened the CLiP Chlamydomonas mutant library under either 35°C or 40°C with and without supplied carbon source using quantitative pooled screens. **(a)** The relative growth rates of Chlamydomonas wild-type cells (CC-5325, CLiP library background strain) at different temperatures with and without supplied organic carbon source. Chlamydomonas cells were grown in photobioreactors (PBRs) in Tris-acetate-phosphate (TAP, with acetate as a carbon source) or Tris-phosphate (TP, no acetate) medium under turbidostatic conditions at different temperatures with a light intensity of 100 µmol photons m^−2^ s^−1^ and constantly bubbling of air. Relative growth rates were calculated based on the cycling of OD_680_, which is proportional to total chlorophyll content in unit of µg chlorophyll mL^−1^. Each temperature treatment was conducted in an individual PBR and lasted two days. Mean ± SD, *n* = 2-5 biological replicates. Statistical analyses were performed using two-tailed t-test assuming unequal variance by comparing with 25°C with the same carbon condition (*, *p* < 0.05, the color and position of asterisks match the treatment conditions). **(b)** Pooled CLiP mutants were grown in PBRs with either TAP or TP medium under different temperatures using the similar cultivation condition as in panel a. Cultures were acclimated for 2 days at 25°C before high temperature treatments. Heat at 35°C (brown lines) lasted for 4 days (TAP) or 5 days (TP). Heat at 40°C lasted for 2 days then recovered at 25°C for 3 days (red) for both TAP and TP conditions. Algal cultures grown at 40°C for longer than two days could not survive. Control cultures were maintained at 25°C for the duration of the experiment (dark blue, dashed lines). Different treatments were conducted in individual PBRs. Samples were collected at the beginning (T1) and the end (T3a or T3b) of the treatment period. T3a was used for the end of TAP-35°C screen, while T3b was used for all other screens due to their slower growth rates than TAP-35°C. **(c)** Schematic of pooled screens utilizing unique internal DNA barcodes of the CLiP mutant library. The Chlamydomonas CLiP mutant library enables high-throughput, quantitative phenotyping in pooled cultures. Three hypothetical mutants are shown, each with a unique DNA barcode sequence (represented by purple, brown, and red bars). CLiP mutants are pooled and grown under the condition as in panel b. Under constant 25°C, three mutants have no change of growth rates, thus no change of barcode abundance. If the brown (or red) mutant is deficient in a process essential for optimal growth under a 35°C (or 40°C) screen condition, it will have reduced cell abundance at the end of the screen. Cell abundance and growth rate of each mutant in pooled cultures are calculated by quantifying DNA barcodes at the start and end of the screen using deep sequencing from pooled mutant DNAs. **(d)** Principal component (PC) analysis of normalized barcode abundance from each sample. The first two PC displayed represent the largest amount of variance explained in the dataset. Closed symbols and solid ovals represent TAP conditions; open symbols and dashed ovals represent TP conditions. Colors represent the different temperature treatment groups (blue: 25°C; brown: 35°C; red: 40°C) and shapes represent sampling time points (T1, T3a, T3b). PVE: percent variance explained.

### Growth rates were quantified consistently

We calculated the growth rates of each mutant using the normalized read counts from DNA barcodes at the beginning and end of the treatment period (see Methods). Because the cell cycle is arrested during 40°C in WT Chlamydomonas cultures (Zhang *et al*. 2022a), growth rates for these conditions were estimated based on the 3-day recovery at 25°C following the 40°C treatment (Figure 1b, Supplementary Dataset 1c, see Methods). Growth rates quantified by barcodes were highly reproducible between two different barcodes from the same mutant (amplified, sequenced, and processed separately, therefore serving as internal controls for pipeline reproducibility) (Figure 2a, b), and relatively reproducible between biological replicates (treatments in separate PBRs) (Figure 2c, d). The relatively lower reproducibility under constant 25°C was partly due to lack of selection pressure. The observed variations in growth rates between biological replicates highlights the need for statistical significance between replicates for downstream analysis (see below). For additional verification, we validated the growth rates of 10 individual CLiP mutants from the pooled mutant screens in both 25°C and 35°C treatment condition in PBRs using monocultures (Figure 2e). Monoculture growth rates were highly consistent with those calculated from pooled mutant screens.

**Figure 2:**
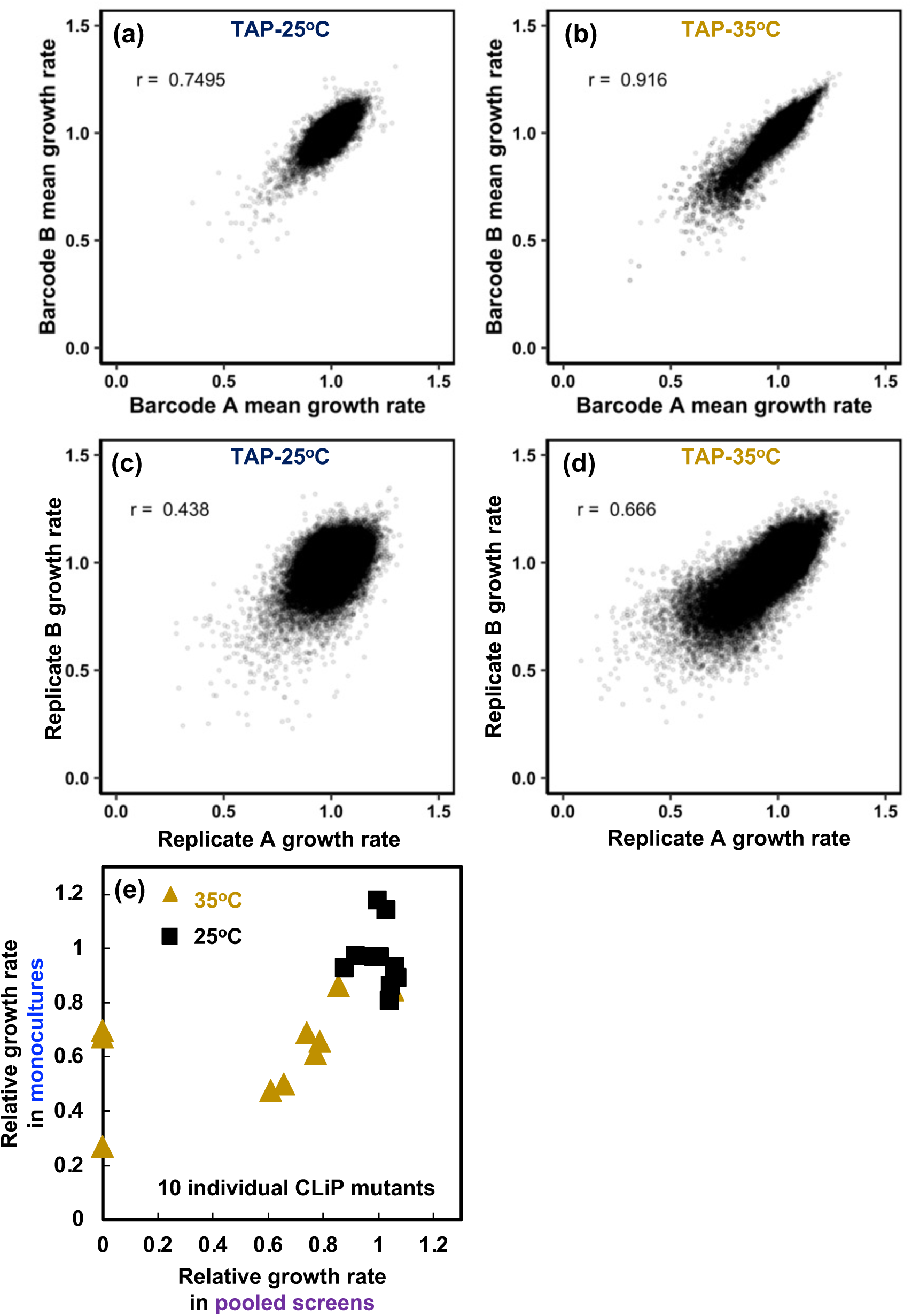
Growth rates calculated from barcode quantification are reproducible. **(a, b)** Growth rates of one mutant with two different barcodes were consistent. Each dot represents one mutant with two different barcodes (from 5’ and 3’ side of the insertional cassette or from 2 different insertion sites). Different barcodes from the same mutants were amplified and sequenced separately. X- and Y-axes are mean normalized growth rates calculated from two different barcodes of the same mutant in the screens of TAP-25°C (**a**) and TAP-35°C (**b**). r: Pearson correlation coefficient. **(c, d)** Growth rates of biological replicates can be calculated consistently. Each dot represents the normalized growth rate of one mutant from two biological replicates in the screens of TAP-25°C (**c**) and TAP-35°C (**d**). Because the screen pressure was present under TAP-35°C but not TAP-25°C, the growth rate plots from the two conditions are different. **(e)** Growth rates calculated from pooled mutant screens were validated with those calculated from monocultures for 10 individual CLiP mutants. X: mean relative growth rates of individual CLiP mutants from TAP-25°C (black) or TAP-35°C (brown) calculated from pooled mutant screens. Y: relative growth rates of individual CLiP mutants calculated from monoculture growth experiments normalized to WT CC-5325 under the same condition. For monoculture experiments, individual CLiP mutants were grown in separate PBRs as in the pooled screens. Two or three biological replicates were conducted for most of the CLiP mutants (see mutant IDs and No. of replicates in Supplemental Dataset 1b). Mutants that were depleted in the heat screens were set to have a growth rate of zero in the pooled screens for the plot.

### Heat-compromised and heat-depleted mutants were identified in the screens

We identified heat-compromised and heat-depleted mutants using growth rates for each of the four screen conditions (Figure 3). Heat-compromised mutants had a mean growth rate at 25°C ≥ 0.95, mean growth rate with high temperature treatments (35°C or 40°C) ≤ 0.8, p < 0.05 and t-value in the 95^th^ percentile (student’s one-sided t-test of unequal variance) (Figure 3, Figure S2, Supplementary Dataset 1d). Heat-depleted mutants had a mean 25°C growth rate ≥ 0.95 but were absent by the end of the screen in at least one high temperature replicate. The aggregated lists of heat-compromised and heat-depleted mutants are referred to as heat-sensitive mutants. Because all heat-sensitive mutants had normal growth rates at 25°C, they had growth defects specifically under high temperature treatments.

**Figure 3:**
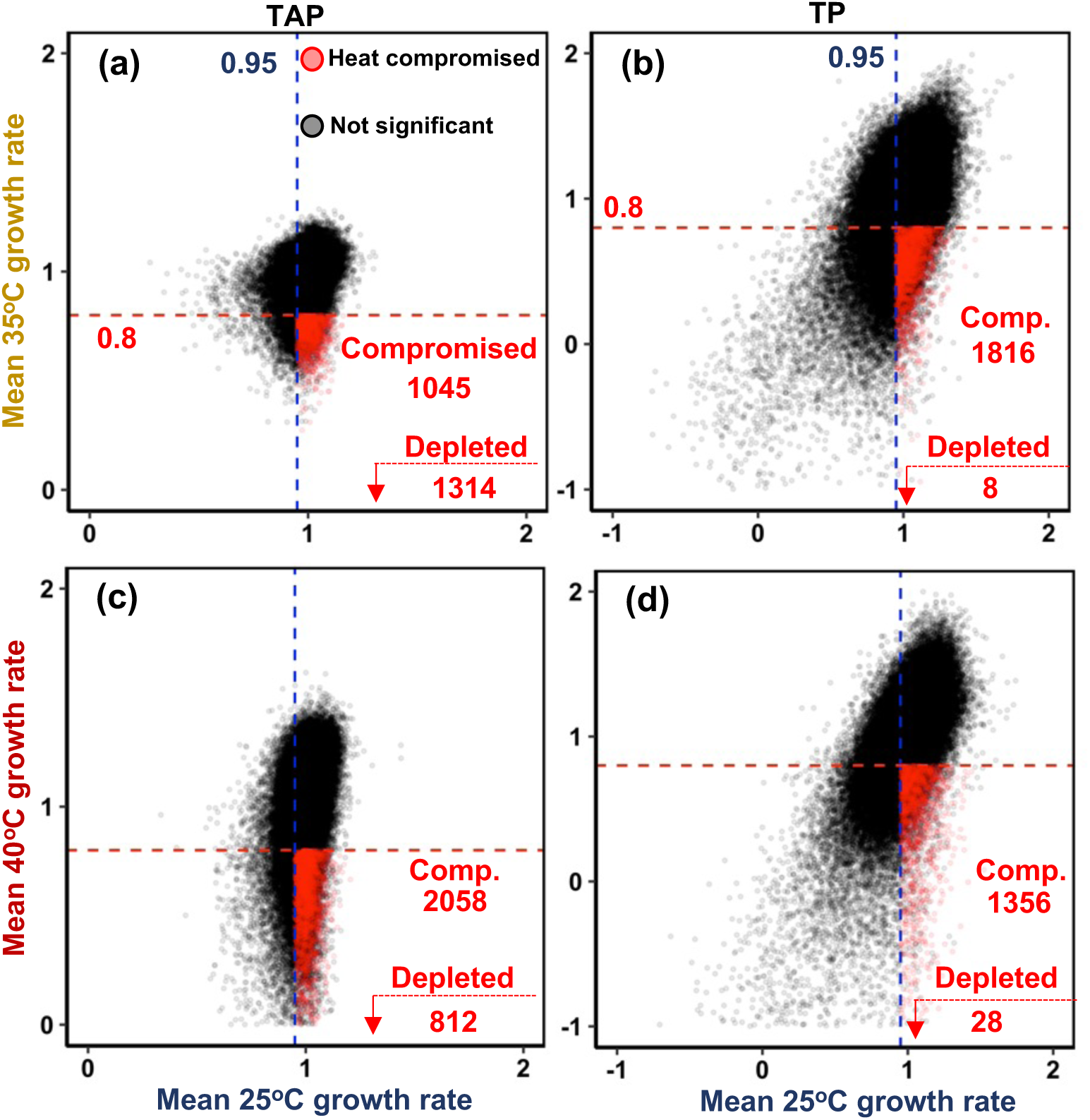
Heat-compromised and heat-depleted mutants were identified from quantitative pooled screens. **(a-d)** Heat-compromised (comp-) mutants from TAP-35°C (**a**), TP-35°C (**b**), TAP-40°C (**c**), TP-40°C (**d**) were defined as those with mean normalized growth rates at 25°C ≥ 0.95, mean normalized growth rates at 35°C or 40°C ≤ 0.8, p-value < 0.05 and t-values in the 95^th^ percentile (student’s one-sided t-test of unequal variance, red dots). Heat-depleted individuals were defined as those that were absent from the 35°C or 40°C pools at the end of the screen but had normal growth at 25°C (≥0.95), indicated in the bottom right corner of each panel.

### We investigated the aggregated features of the heat-sensitive mutants

All CLiP mutants have mapped insertion sites with insertion features and confidence levels (Li *et al*. 2016). The distribution of both insertion features and confidence levels (Figure 4a, b) of heat-sensitive mutants were similar to all CLiP mutants present in our screens. Because each unique DNA barcode is linked to a disrupted gene in a CLiP mutant (Li *et al*. 2016, 2019), we generated a list of genes (4,529) with potential roles in heat tolerance, defined as **<u>H</u>eat <u>T</u>olerance <u>G</u>enes (HTGs)** based on the identified heat-sensitive mutants (Supplemental Dataset 1d). Each HTG has at least one heat-sensitive mutant. While most HTGs had a single mutant allele that was heat sensitive, 5-11% of them had two or more heat-sensitive mutant alleles in a single condition (Figure 4c). Genes with multiple independent heat-sensitive mutant alleles have a high likelihood of being required for thermotolerance.

**Figure 4:**
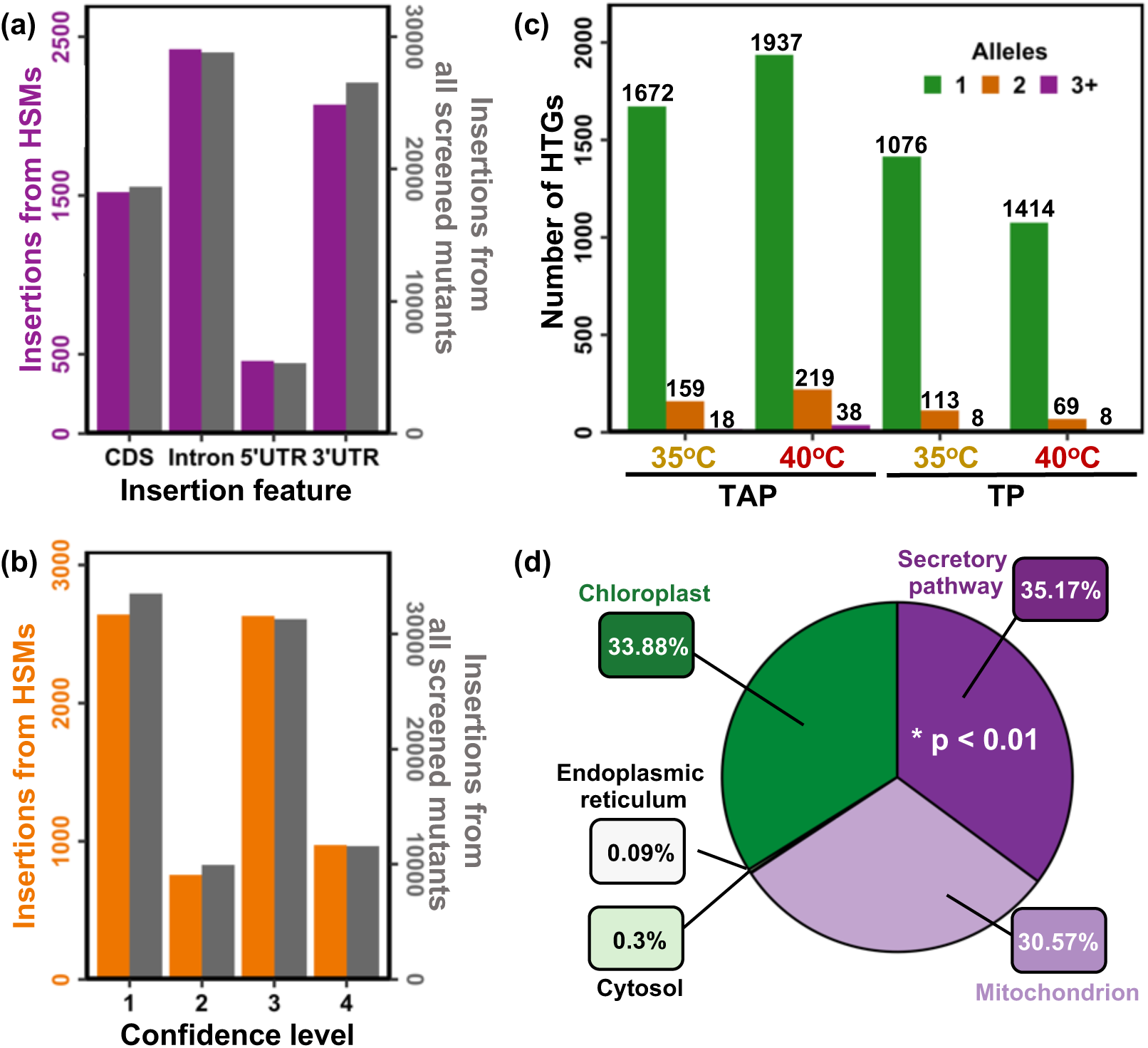
Heat-sensitive mutants disrupted in putative heat tolerance genes (HTGs) were characterized at the genome-wide level. Each CLiP mutant has a mapped insertion site, thus a mapped disrupted gene. We identified putative HTGs from heat-sensitive mutants (HSMs). **(a)** Distribution of insertion features from HSMs (purple) is like that from the CLiP library screened (gray). Insertion feature refers to the integration sites of the transforming cassette relative to gene components, e.g., untranslated regions (UTRs), coding sequence (CDS), or introns. **(b**) Distribution of confidence levels from HSMs (orange) is like that from the CLiP library screened (gray). Insertion confidence refers the likelihood of the mapped insertion sites being correctly identified based on genomic sequences flanking the insertional cassette, with levels 1 and 2 for the highest confidence (95%) of the mapped insertion sites, level 3 for 73% confidence, level 4 for 58% confidence. **(c)** The number of heat-sensitive alleles for each HTG identified from each of the four treatment conditions is shown. Green, orange, and purple bars represent 1, 2, or ≥ 3 heat-sensitive alleles, respectively. **(d)** HTGs identified in any of the four treatment conditions were aggregated and the distribution of predicted subcellular localizations is displayed. Significant enrichment for the secretory pathway was found (*, Fisher’s exact test, p < 0.01). Percentage shown of each localization is the % of HTGs with this predicted subcellular localization out of all HTGs identified with a predicted subcellular localization.

Next, we investigated the predicted subcellular localizations of the aggregated list of HTGs from all conditions using published predictions (Venn & Mühlhaus 2022; Venn *et al*. 2022). We found that significantly more HTGs have a predicted localization in the secretory pathway than expected by random chance (Fisher’s Exact Test, p < 0.01) (Figure 4d, Supplemental Dataset 1d). Of the HTGs with predicted subcellular localizations, 35.17% (818 genes) had proteins predicted to localize in the secretory pathway.

### We identified genes with known and novel roles in thermotolerance

Our HTGs include genes with known roles in thermotolerance (Supplemental Dataset 1e), further supporting the effectiveness of our screen conditions and analysis pipeline. Among these genes were 26 chaperone proteins, which are involved in the proper folding of proteins and are especially important under high temperatures (Schroda *et al*. 2015; Rütgers *et al*. 2017). These include HSP22E HSP70A/H/C, and HSP90A. Additionally, we identified 9 members of the Chlamydomonas carbon concentrating mechanism (CCM) as HTGs. The CCM is responsible for increasing the CO_2_ concentration near the Rubisco active site, thereby increasing the rate of carbon fixation and decreasing the rate of photorespiration, a costly process when Rubisco fixes O_2_ rather than CO_2_ (Mackinder 2018). The CCM is especially important under high temperatures, when the concentration of dissolved CO_2_ to O_2_ decreases in liquid cultures (Blankenship 2014). Furthermore, we identified both forms of Rubisco activase (RCA1/2) as HTGs. RCA is responsible for removing sugar-phosphate inhibitory compounds from the Rubisco catalytic site, which occurs with increased frequency under high temperature and reduces Rubisco activity (Bhat, Thieulin-Pardo, Hartl & Hayer-Hartl 2017; Mueller-Cajar 2017). One of the primary limiting factors for carbon fixation during high temperatures is the activation state of Rubisco (Crafts-Brandner & Salvucci 2002; Perdomo, Capó-Bauçà, Carmo-Silva & Galmés 2017).

Additionally, we identified 2,510 (48%) of HTGs that have no function annotation (Supplemental Dataset 1d), suggesting novel genes with putative roles in thermotolerance. Furthermore, we identified 75 HTGs as transcription factors (Jin *et al*. 2017) (Supplemental Dataset 1f), highlighting the extensive and complex regulation required for coordinating high temperature responses.

### We identified overlapping HTGs between conditions

We next investigated the overlapping HTGs between the four treatment conditions. For this analysis, we only considered those HTGs that were represented by at least one mutant in all four treatment conditions. Consistent with principal component analysis which showed medium treatment had the largest effect on dataset variance (Figure 1d), we found that many HTGs are shared within medium treatments, with 401 genes uniquely having heat-sensitive mutants in TAP conditions and 48 genes uniquely having heat-sensitive genes in TP conditions (Figure 5a, Supplemental Dataset 1d). When comparing temperature treatments, we found 189 HTGs unique to 35°C treatment groups and 172 genes unique to 40°C treatment groups.

**Figure 5:**
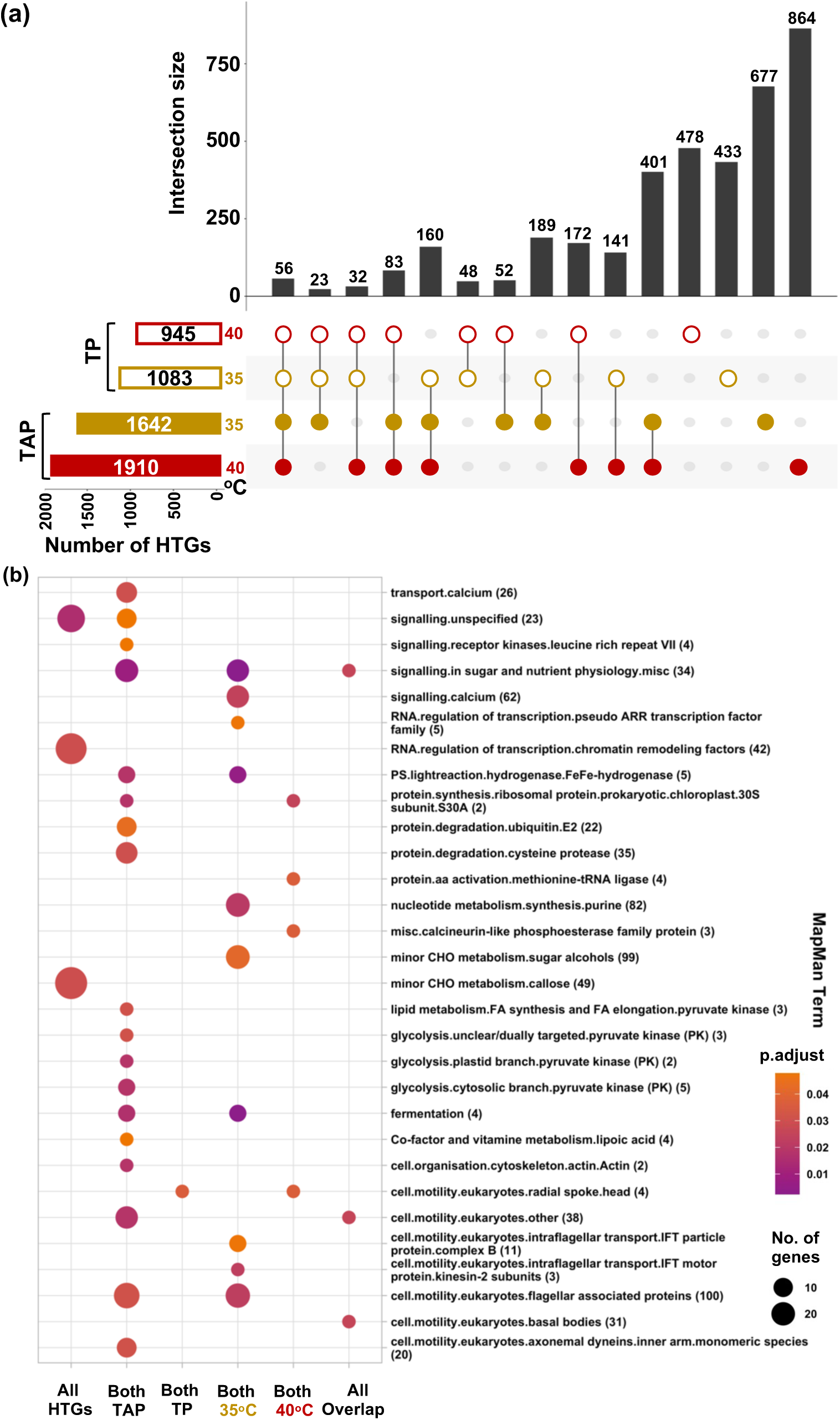
Heat tolerance genes (HTGs) were overlapped between conditions for functional enrichment. **(a)** HTGs overlap between conditions. To compare between conditions, only HTGs represented by at least one mutant in all four conditions were included in this analysis. Horizontal bars represent the total number of HTGs from each condition. Vertical bars and the number on their top represent the number of genes included in each overlap. Circles along x-axis represent the conditions included in each overlap in the vertical bars. TAP, Tris-acetate-phosphate medium; TP, Tris-phosphate medium (no acetate). Filled circles represent TAP conditions and open circles represent TP conditions; circle colors represent the temperature of heat treatments, brown for 35°C and red for 40°C. Genes belong to only a single intersection category and are sorted into the category with the greatest number of intersections. (**b**) MapMan functional enrichment of all HTGs identified (those with at least one heat-sensitive mutant in at least one condition), both TAP conditions, both TP conditions, both 35°C conditions, both 40°C conditions, and all 4 conditions are shown. Only significantly enriched MapMan terms are shown (FDR < 0.05). Size of each circle indicates the number of HTGs in a given MapMan term. Color of the circles represent the FDR adjusted p-value. Numbers in parentheses after each Mapman term is the total number of genes assigned to the given MapMan term and present in the pooled screen dataset.

To better understand the biological functions of HTGs, we performed functional enrichment analysis using MapMan annotations (Figure 5b, Supplemental Dataset 3a-f). In the aggregated list of HTGs from all conditions, we found significantly enriched MapMan terms for signaling, chromatin remodeling factors, and callose. We also investigated the functional enrichment of HTGs in overlapping medium and temperature conditions. For those HTGs found in both TAP conditions (TAP-35°C and TAP-40°C), there were many significantly enriched MapMan terms including calcium transport, iron hydrogenases, protein degradation, lipid metabolism, glycolysis, and cell motility. Interestingly, the enrichment of glycolysis MapMan terms was unique to TAP screens, with the supplied organic carbon source. There was not extensive functional enrichment for HTGs found in both TP conditions, except for cell motility. For HTGs found in both 35°C conditions, calcium signaling, iron hydrogenases, ARR transcription factors, and cell motility were among the significantly enriched MapMan terms. For HTGs found in both 40°C conditions, calcineurin-like phosphoesterases, methionine-tRNA ligases, 30S ribosomal subunit, and cell motility were among the significantly enriched MapMan terms. Finally, we found 56 genes with at least one heat-sensitive mutant in all four treatment conditions, which are significantly enriched for MapMan terms related to nutrient/sugar signaling and cell motility (Figure 5a, Supplemental Dataset 1g, 3f).

### Some HTGs may be also involved in other stresses

We defined HTGs with high-confidence heat-sensitive mutants (HSMs) as those that had either (a) at least two HSMs in one or more treatment conditions or (b) one HSM with a heat-sensitive phenotype in at least two conditions. We next investigated the behavior of these 1,693 HTGs with high-confidence HSMs in other pooled screen conditions conducted by Fauser *et al*. (2022) to see if these HTGs are heat-specific. These authors investigated the behavior of CLiP mutants in approximately 121 different screen conditions (Fauser *et al*. 2022), which we sorted into 16 broad categories (Figure S3). Fauser *et al*. (2022) used two significance thresholds for analysis, FDR < 0.3 and FDR < 0.05. We reported comparisons for the FDR < 0.3 threshold (Figure S3) and included comparisons for both thresholds in Supplemental Dataset 1i & j. For the 16 broad categories, we reported the percentage of genes identified by Fauser *et al*. (2022) with sensitive mutants that are also HTGs in our work. Of our 1,693 HTGs with high-confidence heat-sensitive mutants, we found that 183 also had CLiP mutants there were sensitive to other screen conditions by Fauser *et al*. (2022) (FDR < 0.3), meaning these genes may be also involved in other stresses. Our HTGs with high-confidence HSMs overlapped with 10 of the 39 (FDR < 0.3) and 2 of the 10 (FDR < 0.05) HTGs previously identified by Fauser et al. (2022) (Figure S3, Supplemental Dataset 1i, j), validating our pooled screens. Because both the treatment conditions and analysis pipelines were different between Fauser *et al*. (2022) and ours, we would not expect perfect overlap between HTGs identified. We identified three broad reasons why the remaining 29 out 39 HTGs (FDR < 0.3) Fauser et al. (2022) identified were not included on our HTG list (Supplemental Dataset 1j): (1) some corresponding mutants had no phenotypes in our pooled screen conditions, (2) some mutants had lower growth rates at high temperatures than control, but did not meet the stringent statistical cutoffs we employed, or (3) some mutants had reduced growth rate (<0.95) at control conditions, eliminating them in our analysis pipeline (we required normal growth rates at 25°C ≥ 0.95).

### We investigated HTGs using transcriptomes/proteomes under high temperatures

If a gene is up-regulated at the transcript and/or protein level during high temperatures and disruption of this gene results in a heat sensitive phenotype, this gene likely has an important role in thermotolerance. Thus, we compared our HTGs with our previously published transcriptomes and proteomes in WT Chlamydomonas during 24-hours of moderate (35°C) or acute (40°C) high temperatures followed by 48-hours recovery at 25°C under similar PBR cultivation conditions (Zhang *et al*. 2022a). Our previous RNA-seq identified 3,960 <u>h</u>eat induced <u>g</u>enes (HIGs), up-regulated in at least one time point during the heat treatments of 35°C or 40°C, and 3,503 of them were also present in our pooled screen experiments. We found 1,224 genes overlapping between the HIG and HTG lists, significantly more than expected by random chance (Fisher’s exact test, p < 0.001) (Figure 6a). Because some genes were differentially regulated at the protein level but not the transcript level in our previous data, we were interested in identifying the overlaps between <u>h</u>eat induced <u>p</u>roteins (HIPs, 549 proteins) and HTGs (Figure 6b). We found 111 genes that were both HIPs and HTGs. Unlike the overlap between HTGs and HIGs, the overlap between HTGs and HIPs were significantly de-enriched below what would be expected by random chance (111 observed vs 138 expected). This proteomics dataset was generated by untargeted mass spectrometry, resulting in identification of primarily highly abundant proteins. However, the distribution of HTGs is not over-represented for highly abundant proteins, likely because highly abundant proteins may also be performing important functions under control temperatures. Our HTGs are required to have mutants with normal growth rates at 25°C but reduced growth rates during heat treatments.

**Figure 6:**
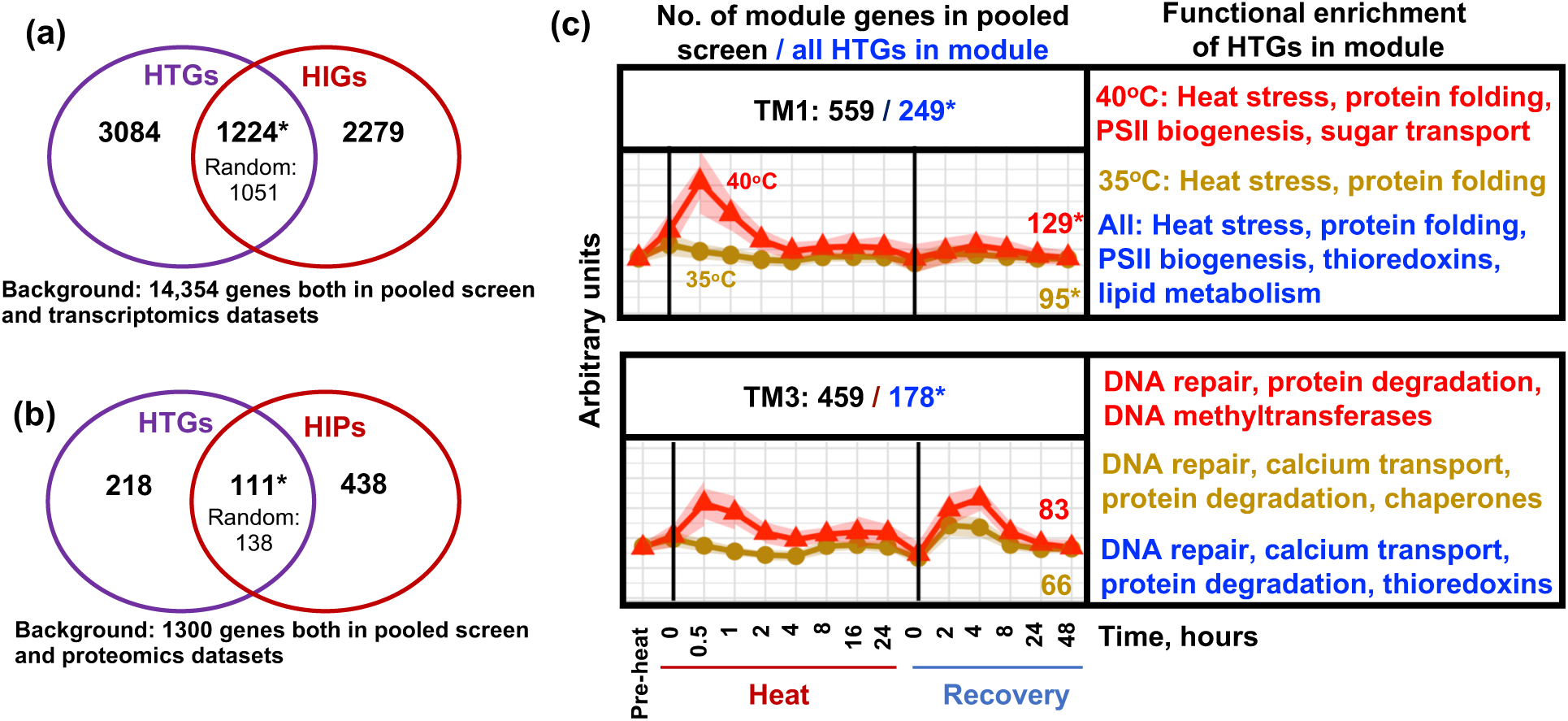
Comparison of heat tolerance genes (HTGs) with transcriptome and proteome data under high temperatures. **(a, b)** Venn diagrams comparing HTGs with heat induced transcripts/proteins (HIGs/HIPs) from 35°C and 40°C treatments of WT cultures in TAP medium (Zhang *et al*. 2022a). Random, expected overlapping numbers based on random chance. *, p < 0.05, Fisher’s Exact Test based on the indicated background size. **(c)** Select weighted correlation network analysis (WGCNA) modules modified from Zhang *et al.,* 2022 that were overrepresented for HTGs above random chance (*, p < 0.05, Fisher’s exact test). TM: transcriptomic module. Consensus transcription patterns from TAP-40°C (red) and TAP-35°C (brown) are shown. Numbers at the top of each module display: the total number (No.) of genes in the given module that were represented by at least one mutant in the pooled screens (before /, black) and the total No. of aggregated HTGs in that module from all four treatment conditions (after /, blue). Numbers at the right of each plot show the No. of HTGs from TAP-40°C (red, top) and TAP-35°C (brown, bottom), respectively. Functional enrichment on right side of each module shows select significantly enriched MapMan terms from TAP-40°C HTGs (top, red), TAP-35°C HTGs (middle, brown) and the aggregated list of HTGs from all four conditions (bottom, blue) that are present in a given module (FDR < 0.05).

Our previous transcriptome analyses also identified common transcriptional patterns during and after high temperatures using weighted correlation network analysis (WGCNA) (Zhang *et al*. 2022a). Interestingly, the HTGs identified in our pooled screens were non-randomly distributed among the WGCNA modules (Figure 6c). Of particular interest is transcriptome module 1 (TM1), which contains 559 genes that are both in TM1 and have at least one mutant in our pooled screens (Supplemental Dataset 1k). These genes had peak expression at the beginning of high temperature and most genes with known roles in thermotolerance (e.g., HSPs) had this transcriptional pattern. Of those 559 genes, 249 were also identified as HTGs in our pooled screens (defined as TM1-HTGs). Furthermore, 129 and 95 of these TM1-HTGs have heat-sensitive mutants in the TAP-40°C and TAP-35°C screen condition, respectively. These subsets are all significantly enriched above random chance (p < 0.05, Fisher’s Exact Test). We performed MapMan functional enrichment analysis on these TM1-HTGs to gain further understanding of their functions (Supplemental Dataset 3g). MapMan terms relating to heat stress, protein folding, PSII biogenesis, thioredoxins, and lipid metabolism were significantly enriched. Half of TM1-HTGs have unknown functions, and these genes are of particular interest for future study. HTGs were also overrepresented in TM3, which contains 459 genes that are present in the pooled screens, and 178 of them were HTGs (p < 0.05, Fisher’s Exact Test) (TM3-HTGs) (Supplemental Dataset 1k). These genes have peaks in expression during both the early high temperature and early recovery periods. The TM3-HTGs were significantly enriched for DNA repair, calcium transport, protein degradation, and thioredoxins (Supplemental Dataset 3h).

### A triangulation approach identified HTGs with increased confidence

To increase the confidence in our putative HTGs, we leveraged our previously published transcriptome and proteome data under high temperatures along with MapMan functional annotations. By triangulating these three datasets, we have narrowed the candidate list of HTGs to those with higher confidence. We required triangulated HTGs to meet two of the three criteria below: (Figure 7a). (**A**) having high-confidence heat-sensitive mutants (HSMs) in our pooled mutant screens (1,693 genes, HTGs that had at least one heat-sensitive mutant in at least two conditions or at least two heat-sensitive mutant alleles across all conditions in our pooled screens). (**B**) The gene was induced by heat in our previously published RNA-seq or proteomics data (4,344 genes) (Zhang *et al*. 2022a). (**C**) The gene has a MapMan annotation related to a pathway known to be involved in heat tolerance or affected by high temperatures (1020 genes, Supplemental Dataset 3j), including heat response (e.g., HSPs), protein folding, lipid metabolism, calcium signaling, photosynthesis, the carbon concentrating mechanism, redox, and other stress responses. Using this triangulation approach, we identified 933 genes that we defined as triangulated HTGs (meet two of the three criteria mentioned above, Supplemental Dataset 1l). Among these, 43 HTGs meet all three of the criteria used, including RCA1/2, the CCM subunits LCI1 and PHC25, heat shock protein HSP70A, and thioredoxin TRXf2. We further refined the list of 933 triangulated HTGs by sorting into HTGs with at least two heat-sensitive mutant alleles (high-confidence triangulated HTG, 386 genes, 41% of triangulated HTGs) and those with a single heat-sensitive mutant allele or present only in the BC overlap category (medium-confidence triangulated HTG, 547 genes, 59% of triangulated HTGs) (Figure 7a).

**Figure 7:**
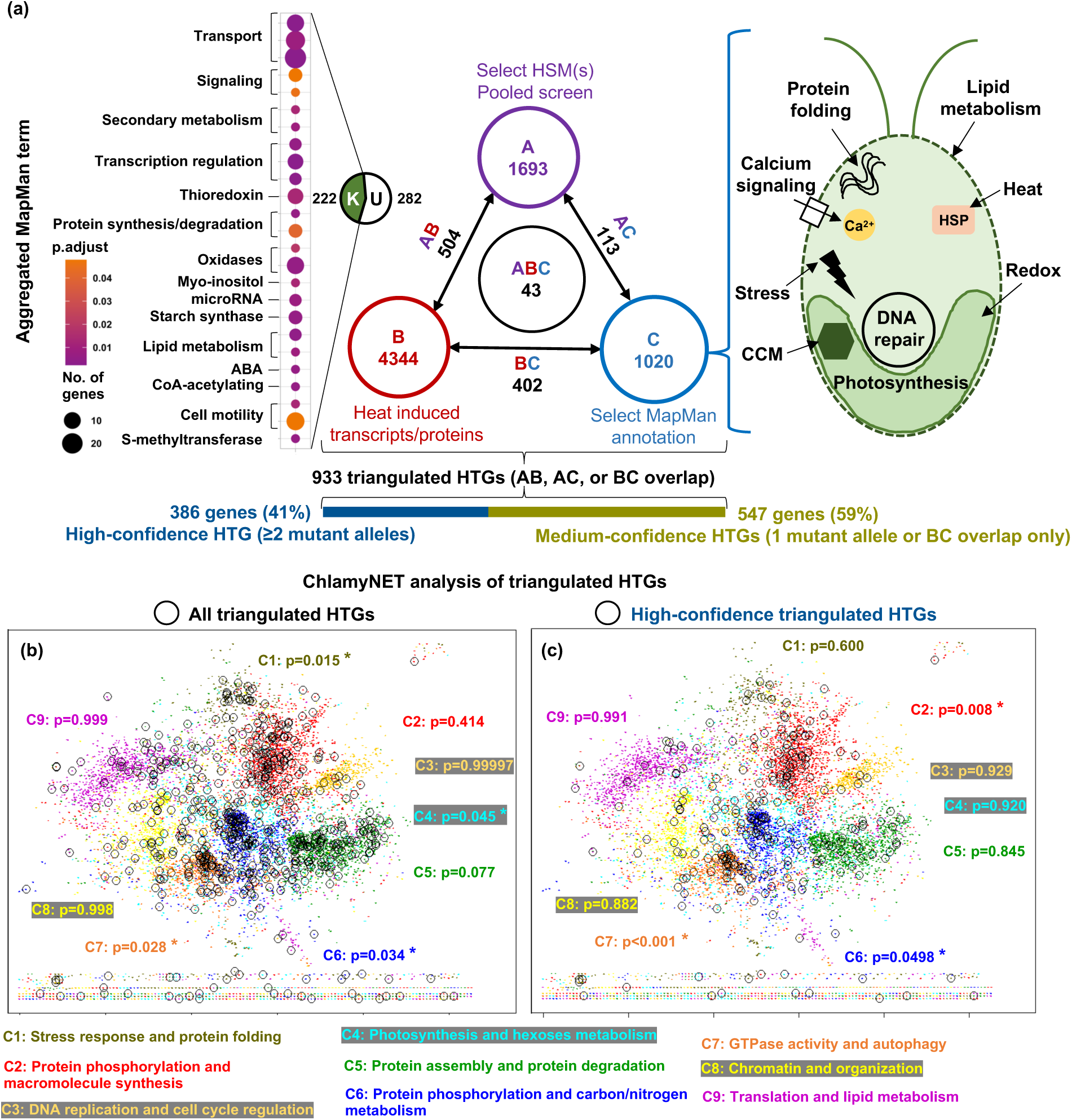
A triangulation approach identified heat tolerance genes (HTGs) with increased confidence. (**a**) Triangulated HTGs were defined based on three criteria: (A): HTGs in our pooled screens that had at least one heat-sensitive mutant (HSMs) in at least two conditions or at least two heat-sensitive mutant alleles in any one condition (defined as high-confidence HSMs); (B): heat inducible during high temperatures based on previously published RNA-seq or proteomic data; (C): MapMan functional annotation in select categories that may be involved in thermotolerance or affected by high temperatures. MapMan functional categories included in this analysis are displayed in cartoon on the right. The number of genes overlapping between any two categories are displayed next to arrows connecting criteria. Of the 504 genes overlapping between criteria A and B, 282 have no MapMan functional annotations (U) and 222 have at least one MapMan functional annotation (K, functional enrichment analysis with abbreviated MapMan terms is displayed). Size of each circle indicates the number of overlapping HTGs in a given MapMan term. Color of the circles represent the adjusted p-value. For full list of MapMan annotations included in this analysis see Supplemental Dataset 3j. HTGs that meet at least two of these three criteria were considered as triangulated HTGs (933 genes), which were further divided into genes with at least 2 heat-sensitive mutant alleles (high-confidence HTGs, 386 genes) and those with a single heat-sensitive mutant allele or present only in the BC overlap (medium-confidence HTGs, 547 genes). (**b, c**) Visualization of triangulated HTGs in the transcription network topology of ChlamyNET, a web-based network tool that was generated based on published transcriptomes in Chlamydomonas (Romero-Campero *et al*. 2016). Nine gene clusters (abbreviated as C) with different function enrichment were identified by ChlamyNET, represented by different colors. The color of a cluster name matches the color of dots in this cluster. The labels of three clusters, C3, C4, C8 with light colors, have grey background to increase the contrast. Each dot represents one Chlamydomonas gene in the network. The genes circled in black are all triangulated HTGs (**b**) or high-confidence triangulated HTGs (**c**). Stars (*) indicate significant enrichment for indicated HTGs within the ChlamyNET clusters (FDR < 0.05, p value for enrichment in each cluster is listed on the figures). The overrepresentation analysis and scatterplot visualization are performed using the R programming language. More information about ChlamyNET analysis can be seen in Supplemental Dataset 1m.

Of particular interest are the 504 genes that meet criteria A and B (heat-sensitive mutants and heat induced transcripts/proteins), because they have two independent experimental datatypes pointing to a role in thermotolerance. Among these, 337 are high-confidence HTGs with at least 2 heat-sensitive mutants. Since this category has no requirement for functional annotation, 282 genes out 504 genes (56%) have no MapMan annotations, suggesting novel players in thermotolerance. The remaining 222 genes were significantly enriched for signaling, transcription regulation, redox, lipid metabolism, starch synthases, and more (FDR < 0.05) (Figure 7a).

There are 113 genes in the intersection between A and C (heat-sensitive mutants and MapMan annotations, 80 of which are high-confidence HTGs with at least 2 heat-sensitive mutants). These genes do not necessarily had increased transcript/protein levels under high temperatures but may encode proteins that have important functions in thermotolerance. For example, some proteins may undergo conformational changes to activate and have no increased abundance to perform their role in thermotolerance.

The intersection between B and C (heat-induced transcripts/proteins and in MapMan annotation) contained 402 genes. There are several reasons why these genes would not be identified in the pooled screen analyses but could still be of great interest. (1) The gene has redundant functionality with another gene, resulting in lack of phenotype in single mutants; (2) the gene may not be represented by knockout mutants in the pooled screen because it is an essential gene, as is the case with the master regulator of heat responses, HSF1; (3) the CLiP mutants have insertions in 3’UTR or introns, thus no observable phenotypes in our pooled screens. Because these genes were not represented by heat-sensitive mutants in this screen, we classified these as medium-confidence triangulated HTGs.

Additionally, we overlayed 933 triangulated HTGs and 386 high-confidence triangulated HTGs with the ChlamyNET dataset, a transcriptional network based on expression patterns of Chlamydomonas under various conditions (Romero-Campero, Perez-Hurtado, Lucas-Reina, Romero & Valverde 2016). Our triangulated HTGs were overrepresented in four ChlamyNET clusters named “C1: stress response and protein folding”, “C4: photosynthesis and hexoses metabolism”, “C6: protein phosphorylation and carbon/nitrogen metabolism**”,** and “C7: GTPase activity and autophagy” (Figure 7b, Supplemental Dataset 1m). Our high-confidence HTGs were overrepresented in three ChlamyNET clusters named “C2: protein phosphorylation and macromolecule synthesis”, “C6: protein phosphorylation and carbon/nitrogen metabolism”, and “C7: GTPase activity and autophagy” (Figure 7c). All triangulated HTGs and high-confidence triangulated HTGs are enriched in both C6 (protein phosphorylation and carbon/nitrogen metabolism) and C7 (GTPase activity and autophagy). The ChlamyNET analysis supports the important roles of our HTGs in thermotolerance.

### Our research in Chlamydomonas can inform orthologous HTGs in land plants

We investigated the conservation of 933 triangulated HTGs (high/medium-confidence levels) in the green lineage. We identified one-to-one, one-to-many, many-to-one, and many-to-many orthologs with eight species using the JGI In.Paranoid dataset (Remm, Storm & Sonnhammer 2001) (Figure 8). Hierarchical clustering identified three distinct classes of conservation for both high-confidence (Figure 8a) and medium-confidence (Figure 8b) triangulated HTGs: (I) low conservation across all species, (II) green-algae specific, and (III) high conservation across all/most species tested. The HTGs identified in class III are of particular interest as our work in Chlamydomonas can be used to infer the function of these HTGs in land plants, providing engineering targets to improve thermotolerance in crops. Of the 933 triangulated HTGs, 173 had a one-to-one orthologous relationship with a gene in the model plant Arabidopsis and 49 of them (28%) were also up-regulated at the transcript level in Arabidopsis during high temperatures (42°C for 7 h) (Balfagón *et al*. 2019) (Supplemental Dataset 1n).

**Figure 8:**
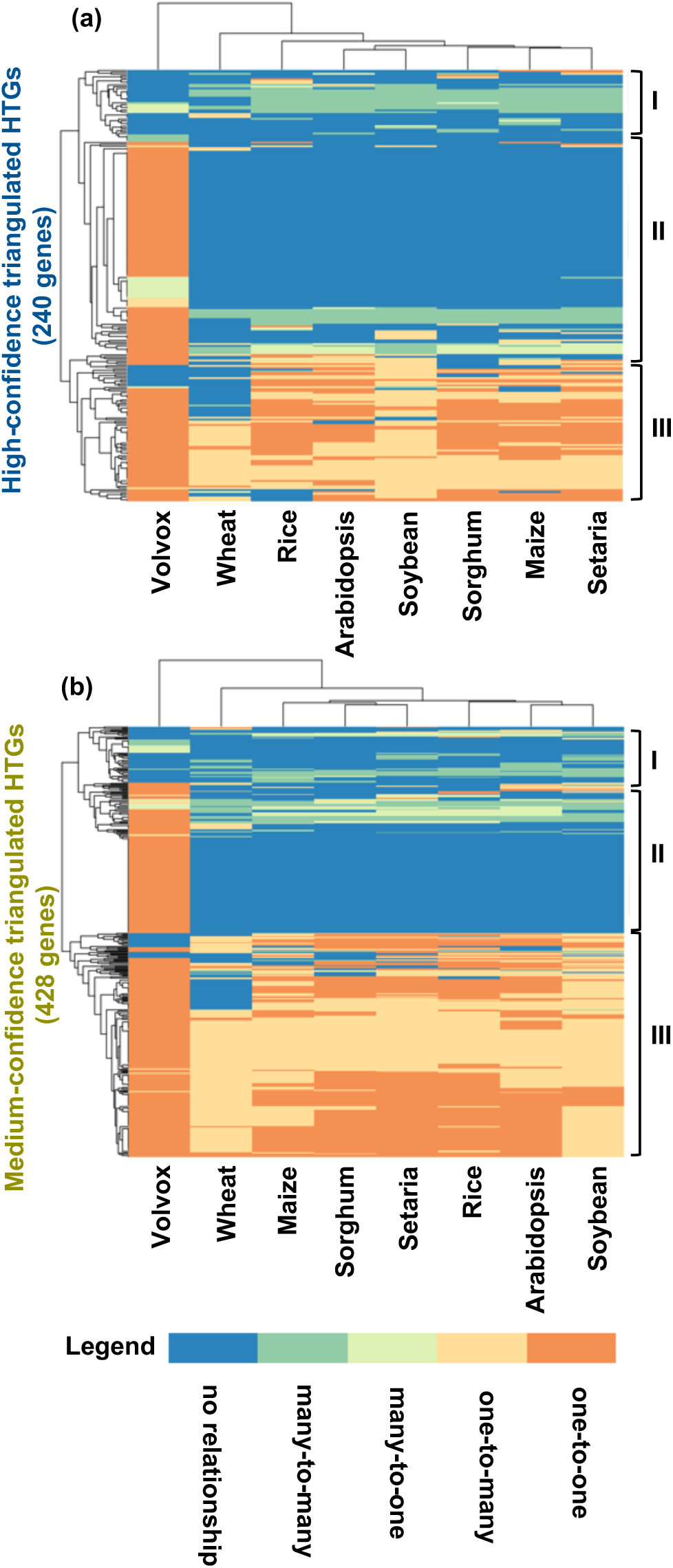
Many Chlamydomonas triangulated heat tolerance genes (HTGs) are conserved throughout the green lineage. Chlamydomonas high-confidence (**a**) and medium-confidence (**b**) triangulated HTGs were classified by their orthology to *Volvox carterii* (Volvox), *Arabidopsis thaliana* (Arabidopsis), *Oryza sativa* (Rice), *Triticum aestivum* (wheat), *Glycine max* (Soybean), *Zea mays* (Maize), *Sorghum bicolor* (Sorghum), and *Setaria viridis* (Setaria) genes using the InParanoid ortholog dataset. One-to-one or one-to-many, one gene in Chlamydomonas corresponds to a single gene or many genes in the species listed. Many-to-one or many-to-many, multiple genes in Chlamydomonas correspond to a single gene or many genes in the species listed. Hierarchical clustering identified three distinct classes of conservation for triangulated HTGs: (I) low conservation across all species, (II) green-algae specific, and (III) high conservation across all/most species tested. Genes were ordered using hierarchical clustering to show similar patterns of conservation across species using the R package pheatmap. Not all HTGs are in the In.Paranoid dataset. Only HTGs in the In.Paranoid dataset with an orthologous relationship with one or more of the species tested are included in the figures.

To demonstrate that our results from Chlamydomonas can be broadly applied in land plants, we investigated the heat sensitivity of Arabidopsis mutants disrupted in orthologs of high-confidence HTGs identified in Chlamydomonas. We required these HTGs to meet the following criteria: (1) there is a one-to-one homolog in Arabidopsis with a Chlamydomonas high-confidence triangulated HTG; (2) the transcript or protein of this HTG is up-regulated during high temperature in Chlamydomonas (Zhang *et al*. 2022a); and (3) the corresponding Arabidopsis transcript is up-regulated during high temperature (Balfagón *et al*. 2019). Among HTGs that meet these criteria, we identified one Chlamydomonas HTG (Cre11.g467752) and its one-to-one ortholog in Arabidopsis (AT4G26240). The Arabidopsis gene is annotated as a putative histone-lysine N-methyltransferase (HMT) at The Arabidopsis Information Resource (TAIR) website, though detailed functional characterization of these two genes is lacking in both organisms.

*CrHMT* was up-regulated at the transcript level in both early heat and early recovery in 40°C, and also during early recovery following 35°C (Figure 9a)(Zhang *et al*. 2022a). *AtHMT* was also up-regulated approximately 1.5-fold following 7 h at 42°C (Figure 9b) (Balfagón *et al*. 2019). Three CLiP mutant alleles in *CrHMT* were heat sensitive in our pooled mutant screens (Figure 9c, d). We obtained two insertional mutant alleles in *AtHMT* (Figure 9e) and tested their heat sensitivity as compared to their WT background, Columbia (Col-0), as well as a well-documented mutant with a heat-sensitive phenotype, *hot-1*, which is mutated in the large heat shock protein HSP101 (Queitsch, Hong, Vierling & Lindquist 2000; Larkindale, Hall, Knight & Vierling 2005; Cha *et al*. 2020). After a 45-minute heat treatment at 41°C followed by 7-day recovery at 25°C, we found *hot1* and *hmt-2* mutants had significantly lower mean plantlet areas as compared to WT (Figure 9f, g). No significant difference was identified for any of the Arabidopsis mutants compared to WT at the control temperature.

**Figure 9:**
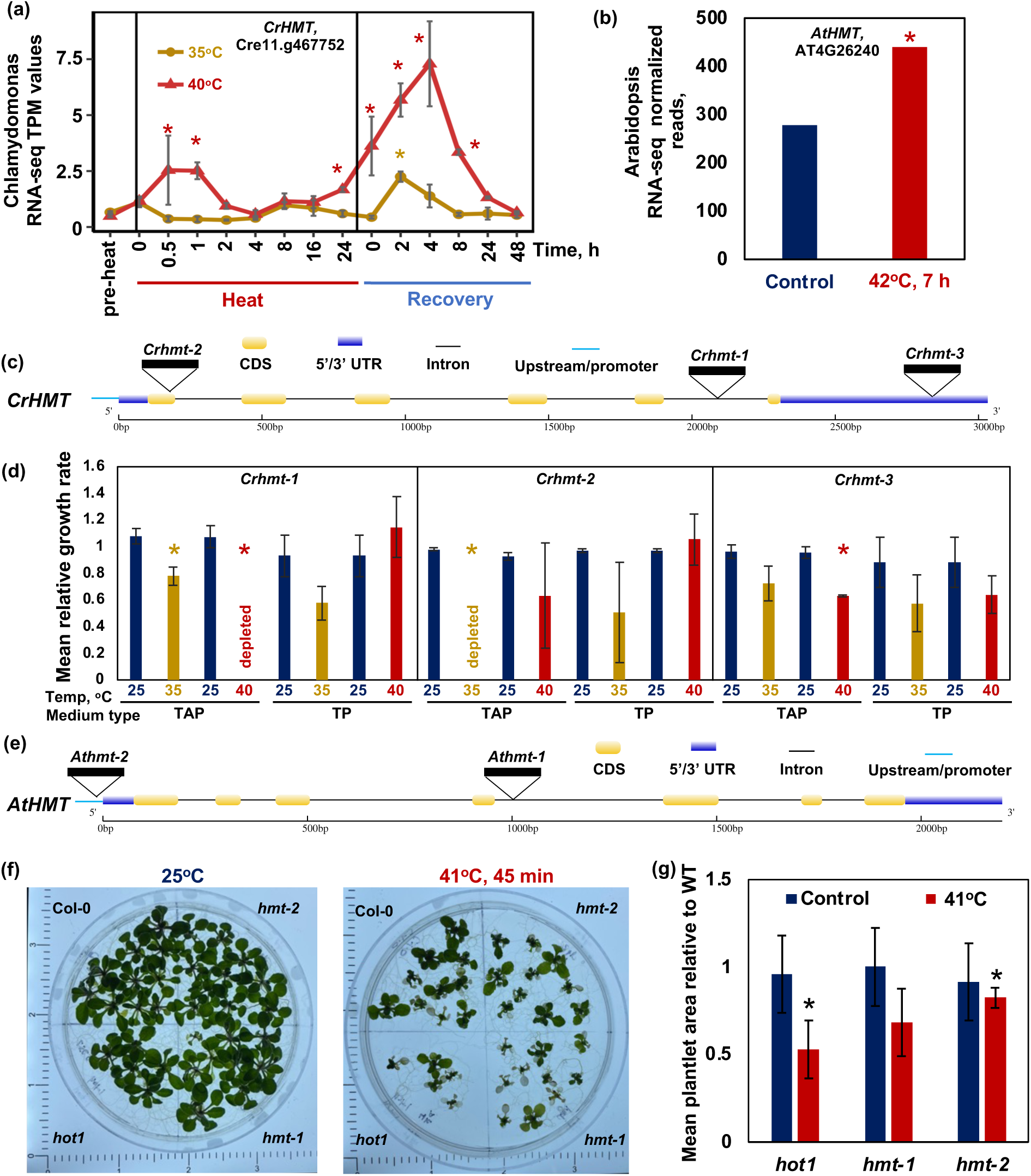
A high-confidence triangulated HTG has heat-sensitive mutant alleles in Chlamydmonas and Arabidopsis. (**a**) Mean transcript per million (TPM) normalized read counts of the Chlamydomonas putative histone-lysine N-methyltransferase (HMT) gene during and after high temperatures of 35°C (brown curve) or 40°C (red curve) (data from Zhang *et. al.* 2022). Mean ± SD, *n*=3. Stars indicate significance in differential expression modeling. (**b**) Normalized RNA-seq read counts of the Arabidopsis one-to-one ortholog (*AtHMT*) of *CrHMT* at control (left) and 42°C (right) (data from Balfagón *et al*. 2019) (**c, d**) Three CLiP alleles of *CrHMT* were heat sensitive in our pooled mutant screens. Stars indicate significance in our pooled screen data. Mean ± SD, *n*=2. TAP, Tris-acetate-phosphate medium; TP, Tris-phosphate medium. (**e**) Two mutant alleles of Arabidopsis *AtHMT*, *Athmt-1* and *Athmt-2* were used for phenotype verification in land plants. (**f**) Representative plate images of 14-day old seedlings from control (25°C) and heat treatment (41°C for 45 minutes followed by 7 days of recovery at 25°C). (**g**) Mean plantlet area relative to wild type (Col-0) of *hmt-1*, *hmt-*2, and *hot1* (positive control, a well-characterized heat-sensitive mutant, deficient in Heat Shock Protein HSP101) in control (left, blue) and heat treatment (right, red) conditions. Plantlet area was quantified using ImageJ and normalized to the total area of Col-0 on the same plate. Mean ± SD, *n*=3-13. *, p < 0.05, student’s one-sided t-test of unequal variance.

## Discussion

We performed genome-wide, quantitative, pooled mutant screens under moderate (35°C) and acute (40°C) high temperatures in medium with or without organic carbon (TAP or TP) using the genome-saturating CLiP mutant library. Through these screens, we identified putative heat tolerance genes (HTGs) in each condition and overlapping conditions. By overlaying these data with previously published transcriptomic and proteomic data during high temperatures and MapMan functional annotations, we report a list of 933 triangulated HTGs (386 and 547 genes at high and medium confidence) in Chlamydomonas, about 50% of which are conserved in other photosynthetic eukaryotes and are prime targets for improving heat tolerance in green algae and land plants. Arabidopsis mutants deficient in the ortholog of a Chlamydomonas high-confidence HTG were heat sensitive, providing evidence that our results in Chlamydomonas can be used to infer thermotolerance genes in land plants.

### Our research expanded the discovery of novel HTGs

Previous pooled screens utilizing the CLiP mutant library employed a data analysis method testing the number of alleles in each gene with a sensitive phenotype against the number of alleles in that gene with a WT-like phenotype (Fisher’s Exact Test) (Li *et al*. 2016, 2019; Fauser *et al*. 2022). Using this method, the authors hope to be confident that the gene of interest is associated with a mutant phenotype in a given condition. However, this method may exclude interesting genes that have many non-phenotypic mutants due to insertions in 3’UTRs or introns, diluting the effect of strong alleles. In our experimental setup, we enabled the use of variance-based statistical methods to identify alleles with consistent heat-sensitive phenotypes. While single mutants showing a heat-sensitive phenotype cannot definitively prove a role of the disrupted genes in thermotolerance, we think these data are valuable for several key reasons: (1) these mutant phenotypes can be combined with other datasets such as multi-omics and functional annotations to add validity to individual mutants; (2) HTGs with strong phenotypes in single CLiP alleles can be used to guide the generation of additional mutants via other mechanisms such as CRISPR (Picariello *et al*. 2020; Dhokane *et al*. 2020) to further characterize the functions of putative HTGs; and (3) these data may be used in the future to characterize the specificity of gene x environment interactions by other groups. Therefore, our triangulation approach comparing our HTGs with heat-induced transcripts/proteins and function annotations increased the confidence of our HTGs with roles in thermotolerance. Nevertheless, to address concerns that heat-sensitive phenotypes may be inaccurately attributed to a gene of interest due to off-target effects, we further refined our triangulated HTGs into those with two or more heat-sensitive alleles (high-confidence, 41%) and those with a single heat-sensitive allele or no CLiP mutants in our screen (medium-confidence, 59%) (Figure 7a). About half of our triangulated HTGs have no or little functional annotation, suggesting novel players with potential roles in thermotolerance.

### We identified core HTGs among different heat treatments

We found 56 HTGs with heat-sensitive mutants in all four heat treatment conditions, representing the list of core HTGs (Supplemental Dataset 1g). Fifteen of these genes had no annotations, representing novel putative HTGs of great interest for future characterization. *CrRCA1* was present in this gene set, highlighting its importance in thermotolerance. Additionally, this gene set contained the PSII assembly/repair factor HCF136. In Arabidopsis, HCF136 is required for the proper insertion of the core reaction center protein, D1, into the PSII complex (Plöchinger, Schwenkert, von Sydow, Schröder & Meurer 2016). High temperatures reduce photosynthesis, increase ROS production, and lead to oxidative stress (Fedyaeva, Stepanov, Lyubushkina, Pobezhimova & Rikhvanov 2014; Babbar, Karpinska, Grover & Foyer 2020; Zhang *et al*. 2022a). The D1 protein is frequently damaged during oxidative stress conditions, resulting in its removal and insertion of a new D1 protein into the reaction center (Järvi, Suorsa & Aro 2015; Theis & Schroda 2016). Without the HCF136 protein, these mutants likely cannot cope with the oxidative stresses caused by high temperatures and perform poorly in pooled screens with heat treatments. Additionally, the list of core HTGs contained several putative signal transduction pathway members, including three adenylate/guanylate cyclases and three 3’-5’ cyclic nucleotide phosphodiesterases. While previous reports have shown that both unfolded proteins (Rütgers *et al*. 2017) and calcium spikes (Königshofer, Tromballa & Löppert 2008; Saidi *et al*. 2009; Wu, Luo, Vignols & Jinn 2012) can lead to the activation of HSFs to induce the heat stress response (Schulz-Raffelt, Lodha & Schroda 2007; Schmollinger, Strenkert & Schroda 2010; Schroda *et al*. 2015), the signaling cascades leading to these responses remain unknown. These six genes are prime candidates for putative members of these signaling cascades.

### HTGs are overrepresented in the secretory pathway

We found that HTGs are overrepresented for proteins localizing to the secretory pathway (Figure 4d). Secretomes are known to contain proteins with a variety of functions including cell wall maintenance, redox homeostasis, and cell-cell communication (Krause, Richter, Knöll & Jürgens 2013; Tanveer, Shaheen, Parveen, Kazi & Ahmad 2014). Recent works across a range of plant species have shown that secretomes vary widely in response to different environmental stresses, including oxidative (Zhou, Bokhari, Dong & Liu 2011; Tanveer *et al*. 2014), drought (Pandey *et al*. 2010; Bhushan *et al*. 2011; Krause *et al*. 2013; Tanveer *et al*. 2014), osmotic (Zhang *et al*. 2009; Song *et al*. 2011; Tanveer *et al*. 2014; Ngara *et al*. 2018), and low temperature (Gupta & Deswal 2012; Tanveer *et al*. 2014) stresses. Furthermore, a recent analysis in Chlamydomonas comparing an evolved salinity-tolerant strain to the parental strain found over 500 differentially accumulated proteins between the two secretomes (Ves-Urai, Krobthong, Thongsuk, Roytrakul & Yokthongwattana 2021). Recently, a high temperature secretome of *Sorghum bicolor* cell cultures identified 31 heat-responsive secreted proteins functioning in protein modification, detoxification, and metabolism (Ngcala, Goche, Brown, Chivasa & Ngara 2020). Though the Chlamydomonas secretome under high temperatures has not yet been explored, we hypothesize that some of HTGs in the secretory pathway may enhance the thermotolerance of Chlamydomonas cells.

### Carbon availability affects the thermotolerance of green algae

The background of the CLiP library, CC-5325, exhibited different growth rates at moderate and acute high temperatures of 35°C and 40°C depending on carbon availability (Figure 1a). Heat at 30°C and 35°C increased the growth of CC-5325 with supplied carbon (TAP medium), but this increase was abolished without supplied carbon (TP medium). These observations led us to hypothesize that the presence of external carbon improves the thermotolerance of Chlamydomonas, at least under moderate high temperatures in our experiments. To understand the molecular mechanisms underpinning this observation, we investigated the functional enrichment of HTGs identified from both the TAP-35°C and TAP-40°C conditions. We found that genes involved in glycolysis, particularly pyruvate kinases, were significantly enriched in this gene set and unique to screen conditions with supplied carbon source (Figure 5b). Glycolysis plays a key role in the breakdown of starch for cellular energy (Johnson & Alric 2012, 2013). Pyruvate kinase catalyzes the final step of glycolysis, converting phosphoenolpyruvate + ADP to pyruvate + ATP (Baud *et al*. 2007; Wulfert, Schilasky & Krueger 2020). Three pyruvate kinases were identified in these significantly enriched MapMan terms: PYK3 (Cre05.g234700), PYK4 (Cre03.g144847), and PYK6 (Cre10.g426292). In plant systems, there are commonly multiple pyruvate kinase isoforms localizing to either the plastid or the cytosol (Plaxton 1996; van Dongen *et al*. 2011). Acetate, present in the TAP medium, is up-taken through the acetate uptake/assimilation pathway then converted to starch via the glyoxylate and gluconeogenesis cycles in Chlamydomonas (Johnson & Alric 2012, 2013).

Indeed, our previous work in WT Chlamydomonas cells showed that most proteins involved in acetate uptake/assimilation and glyoxylate and gluconeogenesis cycles were induced during 35°C heat and starch content increased in 35°C treated cells in TAP medium (Zhang *et al*. 2022a). Increased starch breakdown through the glycolysis cycle and production of cellular energy is likely contributing to the enhanced thermotolerance and increased growth rates of Chlamydomonas in acetate-containing medium during 35°C. The three copies of PYK with heat-sensitive mutants in TAP medium may be important for catalyzing the breakdown of phosphoenolpyruvate and production of ATP under high temperatures. Under low oxygen conditions, allosteric regulation leads to increased activity of PYKs resulting in greater ATP production (van Dongen *et al*. 2011). While the majority of respiratory ATP production typically comes from oxidative phosphorylation in mitochondria, the ATP production from glycolysis can be important under stressful conditions where energy availability is limited (van Dongen *et al*. 2011). A recent proteomics analysis in barley under heat stress also found that utilization of glycolysis as an alternative energy source is important for thermotolerance (Rollins *et al*. 2013). The normal growth phenotypes of the *pky* mutants under control conditions may be explained either by functional redundancy (Chlamydomonas has 6 PYKs), or by stress-specific need for higher enzyme function to increase ATP production for the accumulation of HSPs, maintenance of RCA activity, increased ion channel activities, and other cellular processes under high temperatures.

In comparison to the TAP conditions, there was little functional enrichment for the HTGs in both TP conditions apart from the flagellar radial spoke head, which has a putative role in mechanoregulation of ciliary beating (Grossman-Haham *et al*. 2021). This may be partially explained by the overall smaller overlap between HTGs in both TP conditions as compared to both TAP conditions. While there were 700 (56+83+160+401, No. above vertical bars, Figure 5a) HTGs found in both TAP conditions, representing 42% of the 35°C and 36% of the 40°C HTGs, there were only 159 (56+23+32+48) HTGs found in both TP conditions, representing just 15% of 35°C and 17% of 40°C HTGs. This may be due to the slower growth rates (thus fewer generations) in TP conditions without organic carbon source. Alternatively, this trend may reflect a more diverse set of genes required for tolerance of different high temperatures in the absence of external carbon sources. Future work expanding the length of the screen in TP conditions, thereby increasing the number of generations in the screen, may improve the resolution of HTGs during high temperatures without organic carbon source.

### Different high temperature treatments revealed some non-overlapping HTGs

Next, by comparing the functional enrichment of HTGs in both 35°C treatments to those in both 40°C treatments, we identified functional categories that are differentially required under the different heat treatments. Unlike the comparison between TAP and TP conditions, there were a similar number of HTGs identified in each temperature treatment, with 428 (56+23+160+189) HTGs found in both 35°C conditions and 343 HTGs (56+32+83+172) found in both 40°C conditions (Figure 5a). Within the overlapping 35°C HTGs, calcium signaling was significantly enriched (Figure 5b). Calcium signaling pathways have been frequently implicated in thermotolerance in photosynthetic cells, however the members involved in this pathway are not well understood (Königshofer *et al*. 2008; Saidi *et al*. 2009; Wu *et al*. 2012). Of the 62 genes with annotated roles in calcium signaling represented in this screen, 8 have heat-sensitive mutants in both TAP-35°C and TP-35°C conditions (Figure 5b, Supplemental dataset 3d). These included four putative calmodulin-dependent protein kinases, two putative ATPases, one putative copine protein (Tomsig & Creutz 2002), and one protein with an annotated calmodulin domain but no other known functions. These genes provide novel insight into the putative calcium signaling pathways that are important for the thermotolerance of photosynthetic cells. Of particular interest is FAP39 (Cre02.g145100), a putative cation transporting ATPase localized to the flagella. Some researchers have hypothesized that a heat sensor may be flagellar localized or related to flagellar function in green algae and other organisms (Kamp & Higgins 2011; Sengupta & Garrity 2013; Sekiguchi, Kameda, Kurosawa, Yoshida & Yoshimura 2018), but the mechanisms remain unknown. Furthermore, we identified several cell motility MapMan terms significantly enriched in the 35°C HTGs.

The 40°C HTGs were significantly enriched for calcineurin (CaN) like phosphoesterases. Of the three CaN-like phosphoesterases in this screen, two were identified as HTGs in the 40°C condition (Figure 5b). These calmodulin-dependent serine/threonine phosphoesterases do not have clearly defined roles in Chlamydomonas, but we hypothesize that they are involved in thermotolerance-related signaling pathways. It is possible that these proteins are somehow involved in distinguishing between different intensities of temperature stresses, but the mechanisms of this are unclear. Future work to identify the molecular function of these proteins could provide novel insight into the signaling pathways involved in thermotolerance.

### Our research can be used to improve thermotolerance in land plants

To translate our algal research to land plants, we identified orthologs of triangulated HTGs in key land plants of interest (Figure 8) and about 50% of them are highly conserved in land plants, suggesting we can study the function of these HTGs in Chlamydomonas to infer the function of the orthologous genes in land plants. We highlighted the functionality of our study in Chlamydomonas by testing the heat sensitivity of Arabidopsis mutants deficient in the *AtHMT* gene, which is ortholog to a high-confidence Chlamydomonas HTG. The Arabidopsis mutant *hmt-2* displayed significantly reduced plant size as compared to WT following heat treatments, demonstrating the translatability of our work in Chlamydomonas to land plants. The *AtHMT* gene is annotated as histone-lysine N-methyltransferase, although its detailed function is unknown. Histone post-translational modifications alter nucleosome properties, affect chromatin structure, and impact gene regulation (Crisp, Ganguly, Eichten, Borevitz & Pogson 2016; Ueda & Seki 2020). Histone methylation can either increase or decrease transcription, depending on the amino acid position of the methylation and the number of methyl groups attached to specific histone tails. In Arabidopsis, H3K4me3 (trimethylation of histone H3 at lysine 4) is associated with actively transcribed genes while H3K27me3 is often associated with lowly expressed genes (Oberkofler, Pratx & Bäurle 2021; Friedrich *et al*. 2021; Hu & Du 2022). *CrHMT* has no function annotation. Besides Arabidopsis, *CrHMT* also has one-to-one orthologs in *Setaria viridis, Sorghum bicolor, Volvox carterii, Glycine max,* and *Oryza sativa* (Supplemental dataset 1o). If the annotation of *AtHMT* is correct, it is possible that the epigenetic modifications performed by *AtHMT* and *CrHMT* are required for the regulation of heat responses in photosynthetic cells, which needs function validation in the future. In summary, Chlamydomonas is an excellent model for the study of thermotolerance in photosynthetic cells. Our research in Chlamydomonas advanced understanding of heat responses in photosynthetic cells and identified important engineering targets to improve thermotolerance in both algae and crops.

## Acknowledgements

This research was supported by start-up funding from the Donald Danforth Plant Science Center (DDPSC), sequencing service support from the Department of Energy (DOE) Joint Genome Institute (JGI) Community Science Program (CSP, 503414), and the DOE Biological & Environmental Research (BER) grant (DE-SC0020400) to R.Z. The work (proposal: 10.46936/10.25585/60001126) conducted by the U.S. Department of Energy Joint Genome Institute (https://ror.org/04xm1d337), a DOE Office of Science User Facility, is supported by the Office of Science of the U.S. Department of Energy operated under Contract No. DE-AC02-05CH11231. E.M.M. is supported by the William H. Danforth Plant Sciences Fellowship, DDPSC startup funding to R.Z., and Washington University in St. Louis. J.C. and C. C. was supported by DOE grant (DE-SC0020400) and NIH grant (R01GM093123). J.E was partially supported by the National Science Foundation (NSF) Research Experience for Undergraduates (REU) program at DDPSC, NSF grant DBI-1659812. We are very grateful for Dr. Martin Jonikas for using the CLiP mutant library for our pooled screens. We would like to thank Dr. Martin Jonikas, Dr. Arthur Grossman, Dr. Friedrich Fauser, Dr. Xiaobo Li, Dr. Robert Jinkerson, Dr. Peter Lefebvre, and Weronika Patena for discussion about the CLiP mutant library, Dr. James Umen for feedback, Jeff Berry for helpful suggestions for statistical analyses, Maria Sorkin and Natasha Bilkey for advice regarding Arabidopsis experiments, Dr. Dan Lin for donation of WT Col-0 Arabidopsis seeds, Drs. Saima Sahid, Kaushik Panda, and Benedikt Venn for their helpful suggestions regarding functional enrichment analyses, Riya Patel for helping with photobioreactor maintenance and sample harvesting, Luisa Galhardo for helping with medium preparation, Margarita Ramirez for her preliminary analysis of comparing HTGs with previously published RNA-seq data, and Kaia Luik for helping with some of the ImageJ analysis and manuscript editing.

## Author Contributions

R.Z. supervised the whole project, conducted pooled mutant screens in photobioreactors, and collected cell samples. E.M.M., W.M., and N.Z. prepared samples for barcode extraction and sequencing. DOE JGI CSP team (C.G.D., M. Z., C.P., J.S., R.C.O.), sequenced the DNA barcodes. E.M.M and W.M. developed data analysis pipeline for identification of heat-sensitive mutants. E.M.M. led the project with data analysis, and figure preparation. C.E.B., W.M., and N.Z. completed growth rate validation of individual mutants in monocultures. J.E., E.M.M., and C.E.B. verified the identity of individual mutants used for monoculture phenotyping. C.C. and J.C. performed ChlamyNET analysis. E.M.M. performed heat-sensitivity assay in Arabidopsis. E.M.M. and R.Z. led the writing of the manuscript. C.E.B. and C.C. wrote their corresponding methods. R.Z., E.M.M., J.E., C.E.B., J.C., and C.C. helped revise the manuscript.

## Supplemental Figure Legends

**Figure S1:**
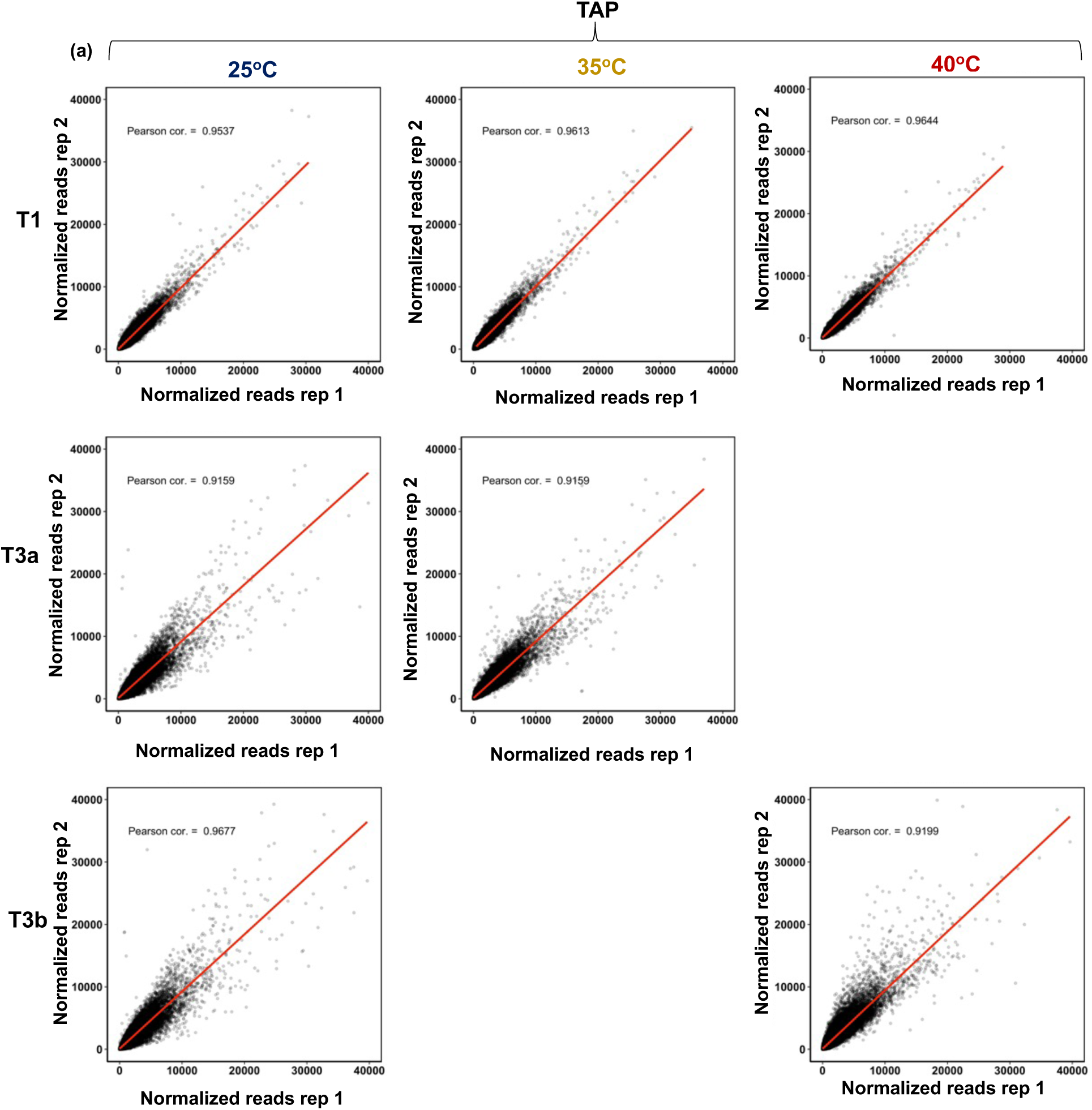
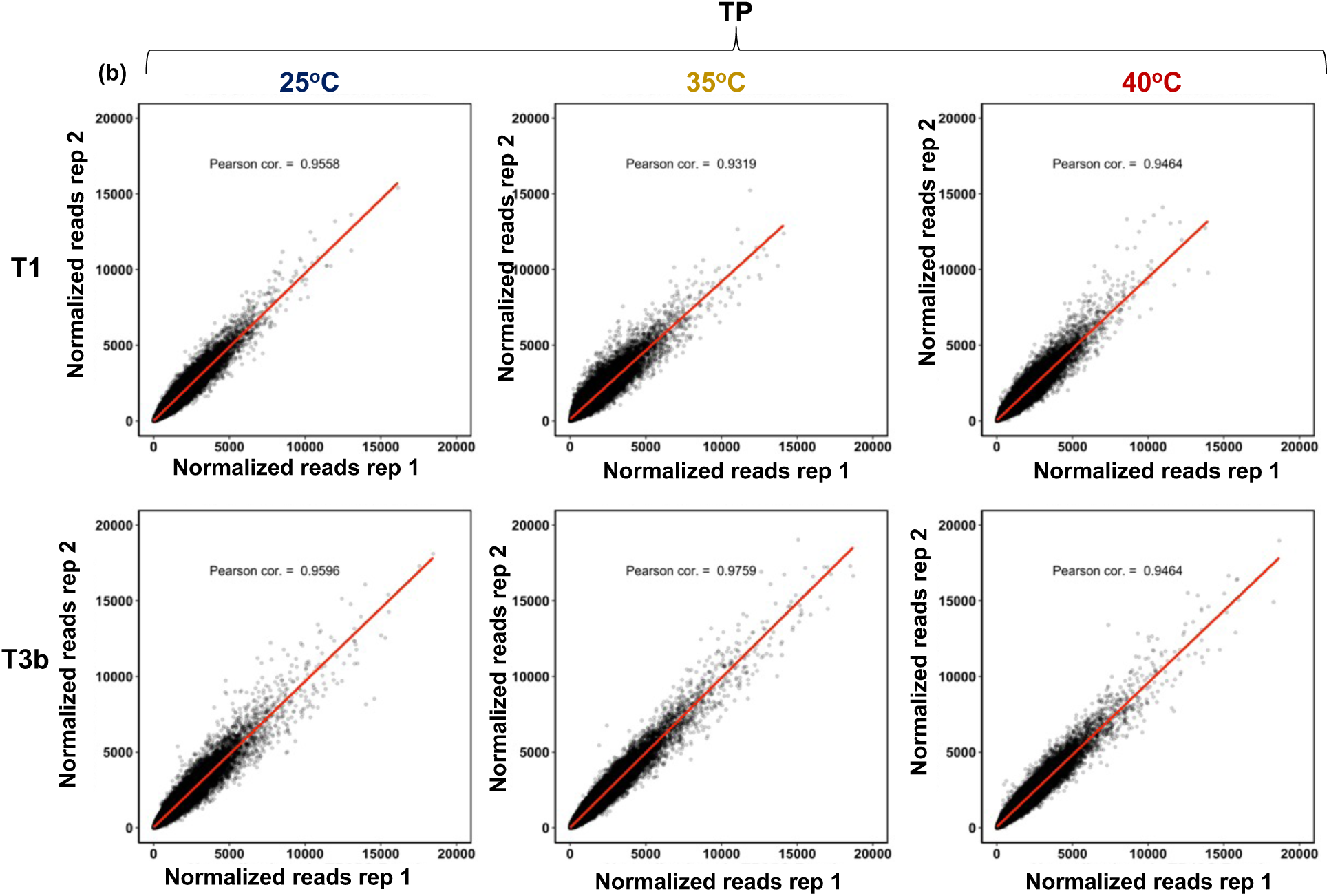
Normalized read counts of biological replicates are highly reproducible. Normalized reads for each barcode in two biological replicates are shown for every condition and time point. Linear model of best fit shown in red line. Pearson correlation coefficient are displayed. Treatment groups are displayed in columns (25°C, 35°C, and 40°C from left to right). Time points are displayed in rows (T1, T3a, T3b from top to bottom. (**a**) All TAP (Tris-acetate-phosphate medium) conditions with supplied carbon source. (**b**) All TP (Tris-phosphate medium) conditions without supplied carbon source.

**Figure S2:**
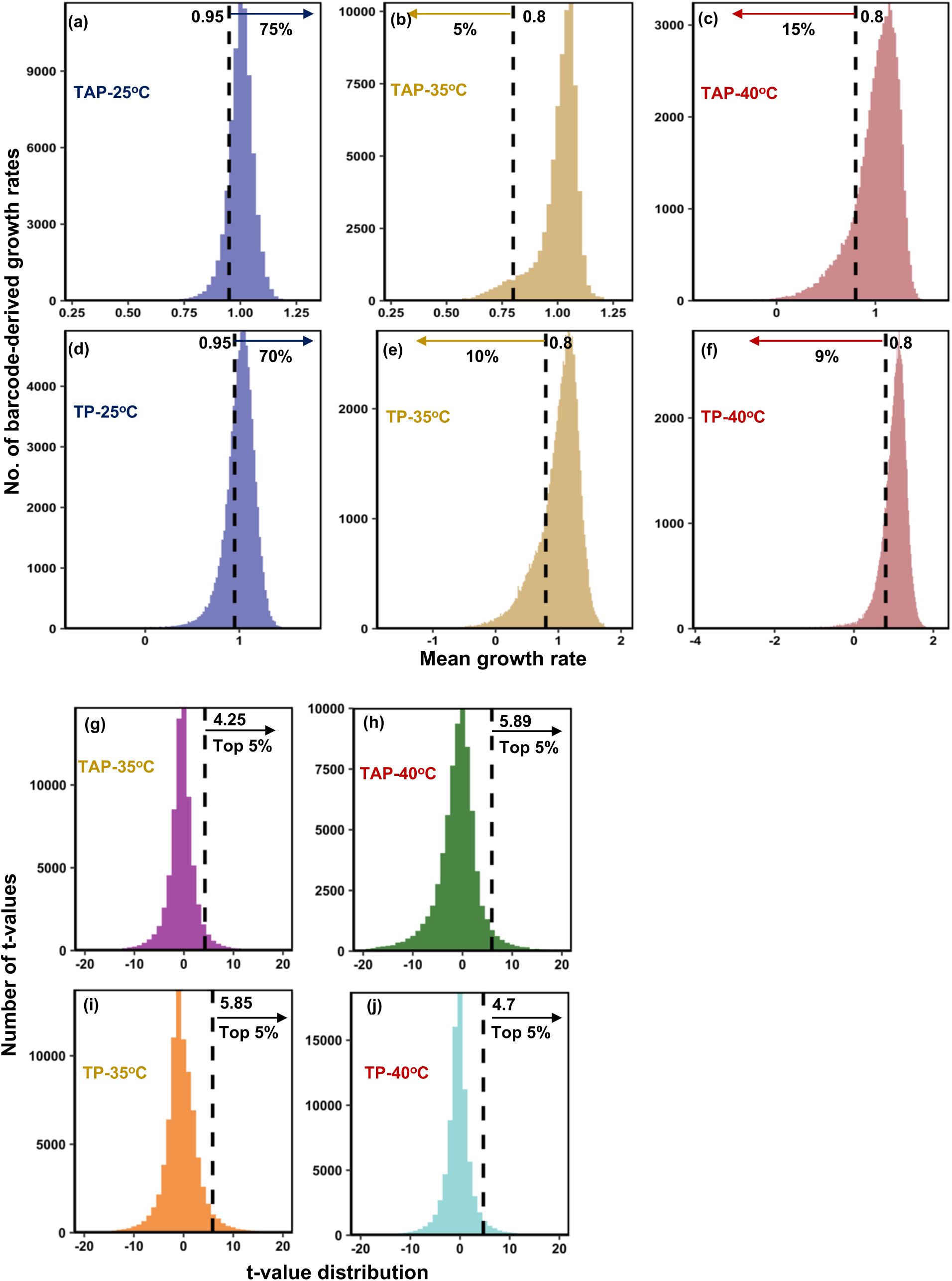
We used statistical cutoffs to identify heat-sensitive mutants. Heat-sensitive mutants are required to have normal growth rates at 25°C (≥ 0.95) and reduced growth rates at 35°C or 40°C (≤ 0.8), p-value < 0.05, and t-values in the 95^th^ percentile (student’s one-sided t-test of unequal variance). (**a-f**) Distribution of mean growth rates of all mutants in each condition. Dashed lines represent growth rates cutoffs, which are displayed at the start of the arrows. Percentage under the arrows quantifies the fraction of barcodes that meet the growth rate cutoffs. (**g-j**) Empirical distributions of t-values for each condition. T-values are based on both the difference between means and the variance between biological replicates. Vertical dashed lines display the 95^th^ percentile mark for each empirical distribution. Numbers above the arrows display the cutoff t-values. Only the top 5% of t-values were considered statistically significant, in addition to p-values < 0.05.

**Figure S3:**
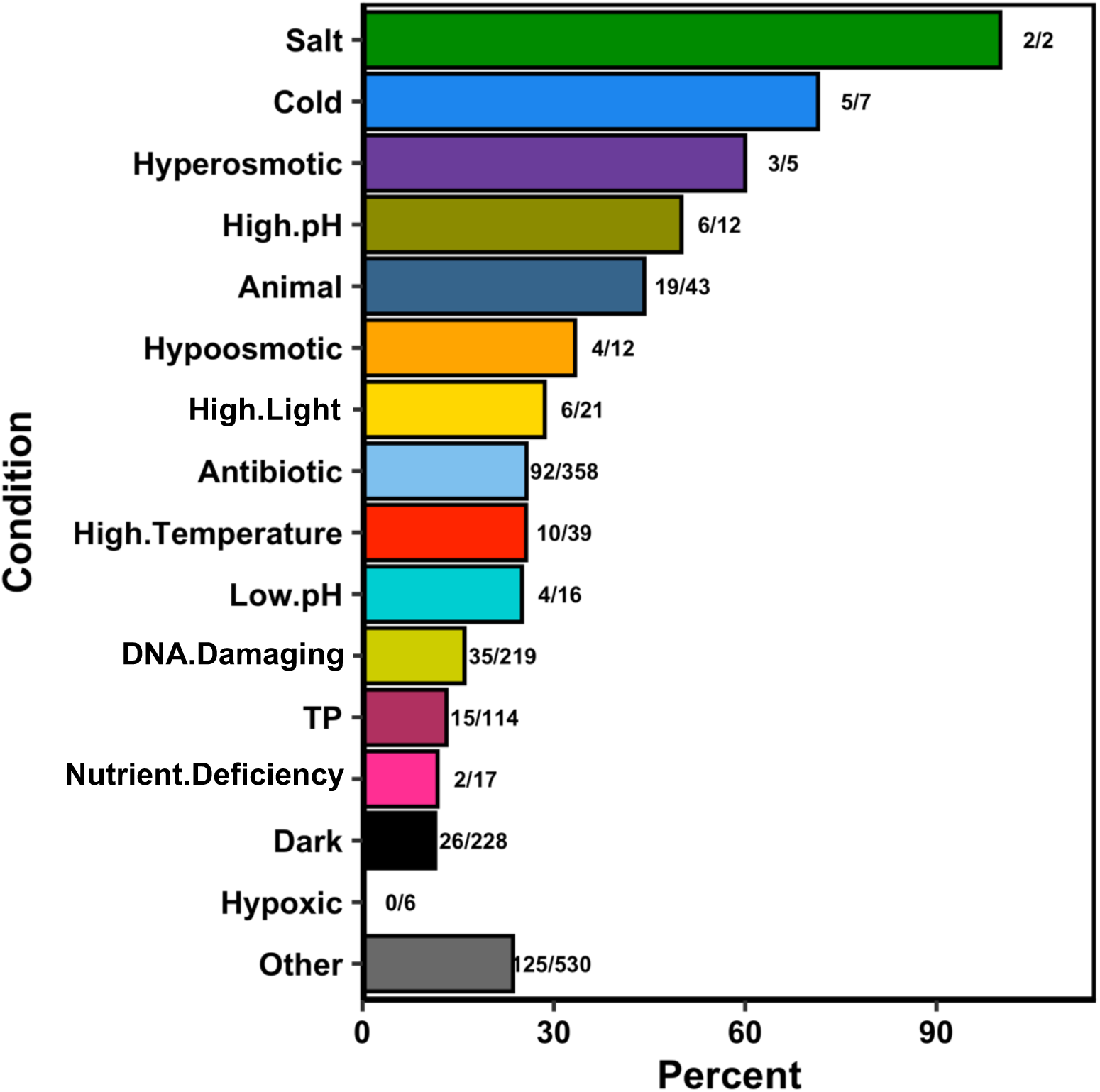
Some heat tolerance genes (HTGs) may be also involved in other stresses. The 1,693 HTGs with high-confidence heat-sensitive mutants (HSMs) identified in our screens were compared with previously published screen data (Fauser et al. 2022) from 121 pooled screen conditions, sorted into 16 broad categories with FDR<0.3. These 1693 HTGs refer to those that have at least one HSM in at least two conditions or at least two HSM alleles in any one condition in our pooled screens. Numbers after horizontal bars are No. of overlapping HTGs with significantly sensitive mutants in that condition (before /) and No. of all genes with significantly sensitive mutants in that condition reported by (Fauser *et al*. 2022) (after /). The ratios of these two numbers are the % presented by the X axis.

**Supplemental Dataset 1:** (**a**) Overview of screen conditions used in this publication and previous pooled screens by Fauser *et al*. (2022). (**b**) List of primers used in this publication. (**c**) Number of generations for each biological replicate used to calculate growth rates of pooled cultures. (**d**) Identification of heat tolerance genes (HTGs) with heat-compromised and heat-depleted mutants in each of all four conditions. (**e**) HTGs with known thermotolerance roles. (**f**) HTGs that are putative transcription factors according to the Plant Transcription Factor Database. (**g**) Core HTGs: HTGs with heat-sensitive mutants in all four conditions. (**h**) List of 1,690 HTGs with high-confidence heat-sensitive mutants. (**i**) Number of genes from the high temperature screens of Fauser *et al*. (2022) that were identified as HTGs with high-confidence heat-sensitive mutants in this study. (**j**) Phenotype of HTGs from Fauser *et al*. (2022) in this study. (**k**) Genes in transcriptional modules 1 and 3 from Zhang *et. al.* 2022 that are represented by at least one mutant in our pooled screens in this study. (**l**) All HTGs that meet triangulation cutoffs (at least 2 of 3 criteria). (**m**) ChamyNET analysis of triangulated HTGs. (**n**) One-to-one orthologs of triangulated HTGs in Arabidopsis. (**o**) For all Chlamydomonas triangulated HTGs, lists the orthologous relationship with species from Figure 8 and the corresponding gene IDs.

**Supplemental Dataset 2:** All pooled screen data calculated from every barcode in this publication.

**Supplemental Dataset 3:** MapMan functional enrichment outputs for all analyses presented in this publication. MapMan functional enrichment for HTGs that are present in: (**a**) at least one heat screen conditions; (**b**) both TAP-35°C and TAP-40°C; (**c**) both TP-35°C and TP-40°C; (**d**) both TAP-35°C and TP-35°C; (**e**) both TAP-40°C and TP-40°C; (**f**) all four heat screen conditions; (**g**) TM1; (**h**) TM3; (**i**) triangulation criteria A and B. (**j**) List of MapMan terms used to generate criterion C of the triangulation approach.

